# Cortical parvalbumin neurons are responsible for homeostatic sleep rebound through CaMKII activation

**DOI:** 10.1101/2023.04.29.537929

**Authors:** Kazuhiro Kon, Koji L. Ode, Tomoyuki Mano, Hiroshi Fujishima, Daisuke Tone, Chika Shimizu, Shinnosuke Shiono, Saori Yada, Junko Yoshida Garçon, Mari Kaneko, Yuta Shinohara, Riina R. Takahashi, Rikuhiro G. Yamada, Shoi Shi, Kenta Sumiyama, Hiroshi Kiyonari, Etsuo A. Susaki, Hiroki R. Ueda

## Abstract

The homeostatic regulation of sleep is characterized by rebound sleep after prolonged wakefulness, but the molecular and cellular mechanisms underlying this regulation are still unknown. We show here that CaMKII-dependent activity control of parvalbumin (PV)-expressing cortical neurons is involved in sleep homeostasis regulation. Prolonged wakefulness enhances cortical PV-neuron activity. Chemogenetic suppression or activation of cortical PV neurons inhibits or induces rebound sleep, implying that rebound sleep is dependent on increased activity of cortical PV neurons. Furthermore, we discovered that CaMKII kinase activity boosts the activity of cortical PV neurons, and that kinase activity is important for homeostatic sleep rebound. We propose that CaMKII-dependent PV-neuron activity represents negative feedback inhibition of cortical neural excitability, which serves as the distributive cortical circuits for sleep homeostatic regulation.

## Main

The sleep-wake cycle is homeostatically regulated (*1*). The wakefulness history is recorded as “sleep need,” which is dissipated during the subsequent sleep. In addition to the brain-wide regulation (*2*), sleep homeostasis is locally regulated in the isocortex (*3*). In humans and rodents, delta power, a sleep need indicator, is locally enhanced in cortical regions that have been more activated during the preceding wakefulness (*4, 5*). Prolonged wakefulness results in “local sleep,” in which cortical neurons locally become silent like in sleep, despite the animal displaying an awake electroencephalogram (EEG) throughout the brain (*6*). These findings suggested that a local and minimal unit of the cortical circuit records their homeostatic sleep need and that the negative feedback system for sleep homeostasis can be found in general cortical circuits.

PV neurons, the most abundant GABAergic interneurons in the isocortex, contribute to feedforward and feedback inhibition in cortical microcircuits (*7, 8*). Thus, we hypothesized that activity control of cortical PV neuron is involved in the regulation of sleep homeostasis. Cortical PV neurons mature throughout postnatal development in terms of distribution, electrophysiological properties, and gene expression (*9*–*13*). Developmental dysfunctions in cortical PV neurons may responsible for the pathogenesis of autism spectrum disorder (ASD) (*14, 15*), and abnormal sleep symptoms are observed in many ASD patients and animal models of ASD (*16*). These findings suggest that the cortical PV-neuron maturation plays a role in physiological sleep-wake regulation. Indeed, sleep architecture and homeostasis are modulated in developmental-stage-dependent manner (*17*–*19*), and cortical PV-neuron activity is modulated by sleep-wake states (*20, 21*). Nonetheless, the role of cortical PV neurons in sleep homeostasis is unknown.

### Sleep architecture and homeostasis are altered along with development

We conducted long-term sleep phenotyping in developing mice after weaning (P21–84) to understand the developmental trajectory of sleep architecture (**Fig. 1A**). There was no significant change in daily sleep duration during the development (**Fig. 1B**). However, both *P*_WS_ and *P*_SW_, the transition probabilities from wakefulness to sleep and sleep to wakefulness (*22*), increased continuously as the developmental stages progressed (**Fig. 1, C** to **D**). The light phase saw a greater increase in *P*_WS_, while the dark phase saw a greater increase in *P*_SW_ (**fig. S1, A** to **E**). Consistent with this, sleep episodes in the dark phase became shorter than that in the light phase during the later developmental stages (**fig. S1F**).

**Fig. 1.**
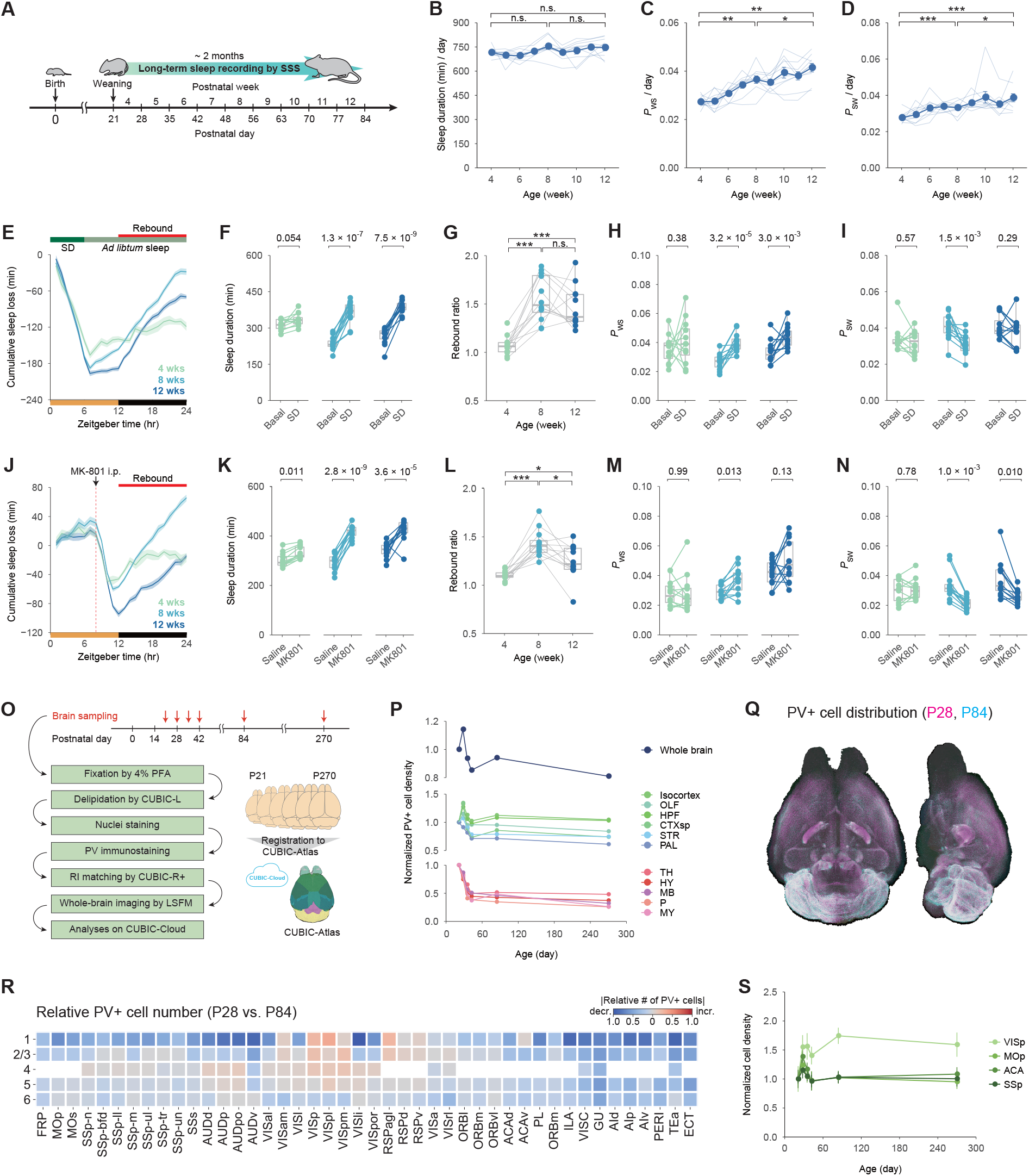
Sleep architecture, sleep homeostasis, and PV-neuron distribution are altered along with development. (**A**) SSS-based long-term sleep phenotyping during post-weaning development (from P21 to P84). (**B, C**, and **D**) Daily sleep duration, *P*_WS_, and *P*_SW_ in developing mice (*n* = 10). The blue points represent the average sleep parameters at each age. Each line (light blue) represents the sleep parameters of a single developing mouse. Welch’s *t*-test with Bonferroni correction was used on mice aged 4, 8, and 12 weeks. (**E**) Cumulative sleep loss (SD – Basal) in mice aged 4, 8, and 12 weeks over 24 hours. (**F, H**, and **I**) Sleep duration, *P*_WS_, and *P*_SW_ on Basal and SD during ZT12–24. At each age, the Welch’s *t*-test was used to compare Basal and SD. (**G**) Sleep rebound ratio (SD / Basal during ZT12–24). Individual mouse dots are connected by gray lines. Between the ages, the Welch’s *t*-test with Bonferroni correction was used. (**J**) Cumulative sleep loss (MK-801 – Saline) in mice aged 4, 8, and 12 weeks over 24 hours. (**K, M**, and **N**) Sleep duration, *P*_WS_, and *P*_SW_ on Saline and MK-801 during ZT12–24. At each age, the Welch’s *t*-test was used to compare Saline and MK-801. (**L**) Sleep rebound ratio (MK-801 / Saline during ZT12– 24). Individual mouse dots are connected by gray lines. Between the ages, Welch’s *t*-test with Bonferroni correction was used. (**O**) Whole-brain analysis of PV+ cell distribution during post-weaning development. P21, P28, P35, P42, P84, and P270 mouse brains were collected (*n* = 6 for each age). Each set of whole-brain images was uploaded to the CUBIC-Cloud and registered to the CUBIC-Atlas brain. (**P**) The mean of PV+ cell density in the whole brain (top), cerebral regions (middle), and brain-stem regions (bottom) across post-weaning development (*n* = 6 for each age). Each point represents the mean value of each age normalized by P21. (**Q**) Representative whole-brain views of PV+ cell distribution from the dorsal and lateral sides at P28 (magenta) and P84 (cyan) ages. (**R**) A region-wise heat map of relative cell number depicting changes in PV+ cell number in the isocortex. P84 and P28 brains were compared, with red and blue colors indicating an increase and decrease, respectively. (**S**) Mean PV+ cell density across post-weaning development in selected cortical regions (*n* = 6 for each age). Each point represents the mean value of each age normalized by that of P21. Error bar: SEM, \**P* < 0.05, \*\**P* < 0.01, \*\*\**P* < 0.001, n.s. = non significance. Brain region acronyms follow the ontology defined by the Allen Brain Atlas.

We further performed a 6-hour sleep deprivation (SD) in individual mice aged 4, 8, and 12 weeks to validate sleep homeostasis in developing mice (**fig. S2A**). The rebound sleep was clearly observed at 8 and 12 weeks old, but less so at 4 weeks old (**Fig. 1, E** to **G**; **fig. S2B**). Wake fragmentation (i.e., increased *P*_WS_) and sleep consolidation (i.e., decreased *P*_SW_) were induced in rebound sleep (**Fig. 1, H** to **I**). Similar to SD, rebound sleep responses were clearer at 8 and 12 weeks old than at 4 weeks old after MK-801 intraperitoneal (i.p.) administration (**Fig. 1, J** to **N**; **fig. S2C**). Long-term sleep phenotyping in developing mice revealed that the sleep-wake pattern and sleep homeostasis mature with post-weaning development.

### Whole-brain distribution of PV neurons is altered along with development

To understand the neural mechanisms underlying the developmental changes in sleep, we focused on PV neurons as a candidate neural basis for the changes. The mouse brains were collected at various stages of post-weaning development, followed by CUBIC-based tissue clearing for a whole-brain analysis of PV-neuron distribution (*23, 24*) (**Fig. 1O**). The averaged PV-positive (PV+) cell density in the whole brain was increased from P21 to P28 and gradually decreased with subsequent developmental stages (**Fig. 1P**, top panel). Similar trends were observed in many cerebral regions (**Fig. 1P**, middle panel), while the PV+ cell density decreased monotonically in all brain-stem regions (**Fig. 1P**, bottom panel). These changes could be attributed to PV+ cell loss rather than a decrease in PV expression in individual PV+ cells (**fig. S3, A** to **B**).

A detailed comparison of PV-neuron distribution in the P28 and P84 brains revealed subregion-specific changes in PV+ cell density (**Fig. 1, Q** to **R**; **fig. S3C**). Except for the visual (VIS) cortex, the PV+ cell density in P84 was lower than in P28 in almost all areas of isocortex (**Fig. 1R**). PV+ cell density in the somatomotor (MO) and anterior cingulate (ACA) cortex gradually decreased beginning at P28, whereas it increased or stabilized during the same period in the VIS cortex (**Fig. 1S**). The PV+ cell density in some cortical regions, such as the somatosensory (SS) cortex, changed little during development (**Fig. 1S**).

### Cortical PV neurons are activated upon the prolonged wakefulness

We investigated PV-neuron activity during SD to understand the physiological role of PV neurons in sleep homeostasis. After a 6-hour SD, mouse brains were collected for whole-brain immunostaining of PV and c-Fos proteins to investigate PV-neuron activity upon SD (**Fig. 2A**). A comparison of *ad libitum* sleep (Slp) and SD brains revealed that SD increased c-Fos-positive (c-Fos+) cell density in many brain regions (**Fig. 2B**). There was also an increase in c-Fos+ cell density among the PV+ cell population (i.e., c-Fos+PV+ cells), particularly in cortical regions (**Fig. 2C**). We also discovered that the SD increased the density of PV+ cell in the isocortex to some extent (**Fig. 2D**). The population of c-Fos+PV+ cells was higher in SD brains than in Slp brains in the isocortex (**Fig. 2, E** to **F**; **fig. S4, A** to **B**). Cortical c-Fos+PV+ cells were found to be increased in MK-801-treated brains (**Fig. 2, G** to **H**; **fig. S4, C** to **G**). These findings indicate that prolonged wakefulness increases cortical PV-neuron activity.

**Fig. 2.**
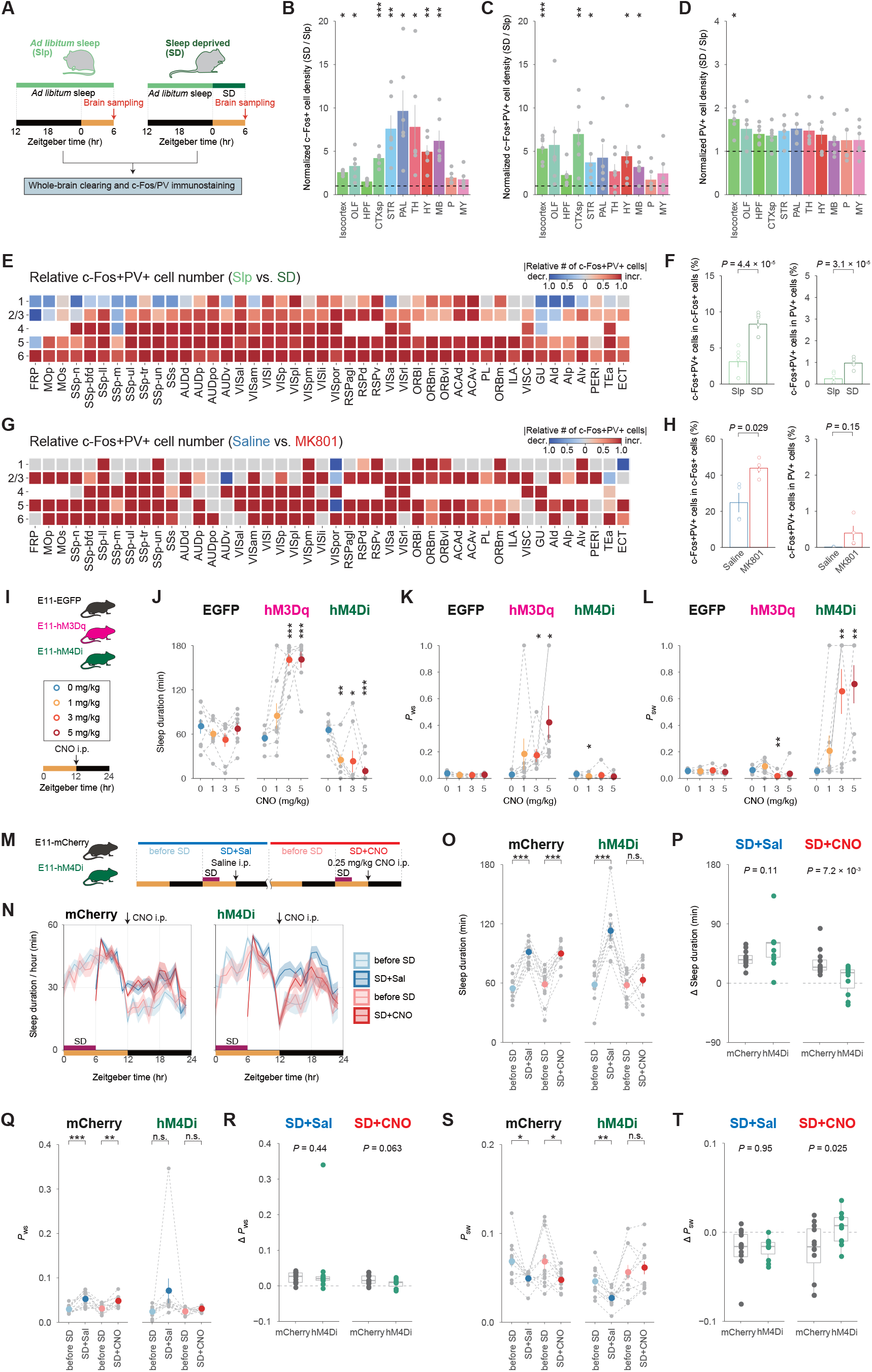
PV-enhancer-targeted (E11) neurons are necessary and sufficient for rebound sleep. Experiment design for whole-brain analysis of c-Fos+ and/or PV+ cell distribution sleep-deprived condition. At ZT6, the brains of control (Slp) and sleep-deprived (SD) mice were collected (*n* = 6 for each group). Each set of whole-brain images was uploaded to the CUBIC-Cloud and registered to the CUBIC-Atlas brain. (**B, C**, and **D**) Normalized cell density (SD / Slp) of c-Fos+ cells, c-Fos+PV+ cells, and PV+ cells in major brain regions (*n* = 6 for each group). Each gray dot represents an individual SD brain normalized by the mean of Slp brains (mean = 1, black-dashed lines), and the colored bars represent the mean of SD brains in each brain region. The Welch’s *t*-test was applied to the normalized value of Slp brains. (**E** and **G**) A region-wise heat map of relative cell number depicting changes in c-Fos+PV+ cell number in the isocortex. SD brains were compared to Slp brains, and MK-801-injected mouse (MK-801) brains were compared to control (Saline) brains, with red and blue colors indicating an increase and decrease, respectively. (**F** and **H**) Rate of double-positive (c-Fos+PV+) cells in c-Fos+ cells (left) or PV+ cells (right) in the isocortex under SD or MK-801 administration. The groups were compared using Welch’s *t*-test. (**I**) Illustration of the CNO injection. Each mouse received i.p. injection in turn: 0 (blue), 1 (gold), 3 (red), or 5 (dark red) mg/kg CNO injection at ZT12, with at least 2 days between injections. (**J, K**, and **L**) Sleep duration, *P*_WS_, and *P*_SW_ on the days with CNO injection during ZT12–15 (*n* = 8 for each group). Each gray dot represents an individual mouse, while colored dots represent the mean in each condition. Individual’s mouse dots are connected by gray-dashed lines. Within the groups, Welch’s *t*-test were run against the 0 mg/kg CNO condition. (**M**) Schematic diagram of CNO injection after sleep deprivation (SD). Every mouse received i.p. injection after a 6-hour SD during ZT0–6. At ZT12, 0 (blue, SD+Sal) or 0.25 (red, SD+CNO) mg/kg CNO injection was given. One day before the scheduled day with SD and i.p. injections are labeled “before SD.” (**N**) Sleep duration over 24 hours in E11-mCherry (*n* = 12) and E11-hM4Di (*n* = 11) mice on SD+Sal (blue), SD+CNO (red), and “before SD” (light blue and salmon) days. (**O, Q**, and **S**) Sleep duration, *P*_WS_, and *P*_SW_ on the days shown in the panel (N) during ZT12–15. Each gray dot represents an individual mouse, while the colored dots represent the mean in each condition. Individual mouse dots are connected by gray-dashed lines. Welch’s *t*-test was used to compare the day with SD to the day “before SD.” (**P, R**, and **T**) The differences (Δ) in sleep duration, *P*_WS_, and *P*_SW_ between SD day during and the “before SD” day during ZT12–15. Welch’s *t*-test was used to compare the groups in the SD+Sal or SD+CNO conditions. Error bar: SEM, \**P* < 0.05, \*\**P* < 0.01, \*\*\**P* < 0.001, n.s. = non significance. Brain region acronyms follow the ontology defined by the Allen Brain Atlas.

### PV-enhancer-targeted (E11) neurons are necessary and sufficient for rebound sleep

Next, we aimed to test the influence of PV-neuron activity on sleep architecture and homeostasis. Cortical PV neurons were targeted using E11, a PV-neuron selective enhancer derived from the *Pvalb* region (*25*), packaged in adeno-associated virus (AAV) vector (**fig. S5A**). Nuclear-localized mCherry signals under the E11 enhancer (E11-H2B-mCherry) were primarily detected in cerebral regions including the isocortex, and more than 80% of E11-labeled cells in the isocortex were PV+ cells (**fig. S5, B** to **E**). The cells targeted by this system were termed as E11 neurons after that.

We expressed a tetanus-toxin light-chain (TeLC) protein, which blocks synaptic transmission, in E11 neurons. TeLC expression in E11 neurons resulted in a negligible change in sleep duration but a significant increase in *P*_WS_ and *P*_SW_ (**fig. S5, F** to **H**). We also conducted chemogenetic manipulations of E11-neuron activity (**Fig. 2I**). Mice expressing hM3Dq (E11-hM3Dq mice) had an increase in sleep duration after clozapine-*N*-oxide (CNO) administration (**Fig. 2J**; **fig. S5, I** to **J**). In the E11-hM3Dq-induced sleep state, increased *P*_WS_ and decreased *P*_SW_ were observed (**Fig. 2, K** to **L**; **fig. S5, K** to **N**), similar to the rebound phase after SD (**Fig. 1, E** to **G**). Chemogenetic suppression, on the other hand, resulted in an acute reduction in sleep duration with decreased *P*_WS_ and increased *P*_SW_ in mice expressing hM4Di (E11-hM4Di mice) (**Fig. 2, J** to **L**; **fig. S5, I** to **N**).

We hypothesized that E11-neuron activity is required for rebound sleep because acute enhancement of E11-neuron activity induced a rebound-sleep-like state (**Fig. 2, J** to **L**; **fig. S5, I** to **N**). To test this, we used chemogenetic inhibition of E11 neurons after SD (**Fig. 2M**). We used a lower CNO dosage that had little effect on the normal sleep-wake cycle (**fig. S6, A** to **G**). Even in this condition, E11-hM4Di mice had impaired rebound sleep after CNO administration (E11-hM4Di mice: SD+CNO), whereas normal rebound sleep was observed in the absence of CNO administration (E11-hM4Di mice: SD+Sal) (**Fig. 2, N** to **P**). Mice only expressing mCherry (E11-mCherry mice) exhibited normal rebound sleep with or without CNO administration (E11-mCherry mice: SD+Sal or SD+CNO) (**Fig. 2, N** to **P**). Rebound responses in *P*_WS_ and *P*_SW_ were also reduced in E11-hM4Di mice after CNO administration (**Fig. 2, Q** to **T**). These findings suggested that rebound sleep requires increased E11-neuron activity.

### Inhibition of CaMKII activity in E11 neurons impairs sleep homeostasis

We then concentrated on CaMKII as a key molecule since the phosphorylation states of CaMKII and its potential substrates in forebrain synapses change with sleep-wake states (*26*–*28*), and animals with CaMKIIα/β genetic modification had abnormal sleep architecture (*29, 30*). Using single-cell RNA sequencing (scRNA-seq) data (*31*), we confirmed that all CaMKII isoforms (*Camk2a, Camk2b, Camk2g*, and *Camk2d*) were expressed in cortical PV neurons (**fig. S7, A** to **D**). Furthermore, using other scRNA-seq data (*32*), we found that *Camk2a* expression level and positive-cell rate in cortical PV neurons were higher in the adults compared to adolescents (**fig. S7, E** to **H**).

To see if CaMKII within PV neurons regulates sleep architecture and homeostasis based on its kinase activity, we inhibited the endogenous CaMKII activity in E11 neurons with CN19o, an optimized CaMKII inhibitory peptide (*33*) (**Fig. 3A**). Mice expressing CN19o under the E11 enhancer (E11-CN19o mice) showed no significant difference in sleep duration when compared to mice expressing CN19scr (E11-CN19scr mice) (**fig. S8A**), the CN19o scrambled sequence (**Fig. 3A**). However, the E11-CN19o mice had higher *P*_WS_ and *P*_SW_ than the E11-CN19scr mice (**fig. S8, B** to **C**), similar to E11-neuron chronic silencing (**fig. S5, F** to **H**).

**Fig. 3.**
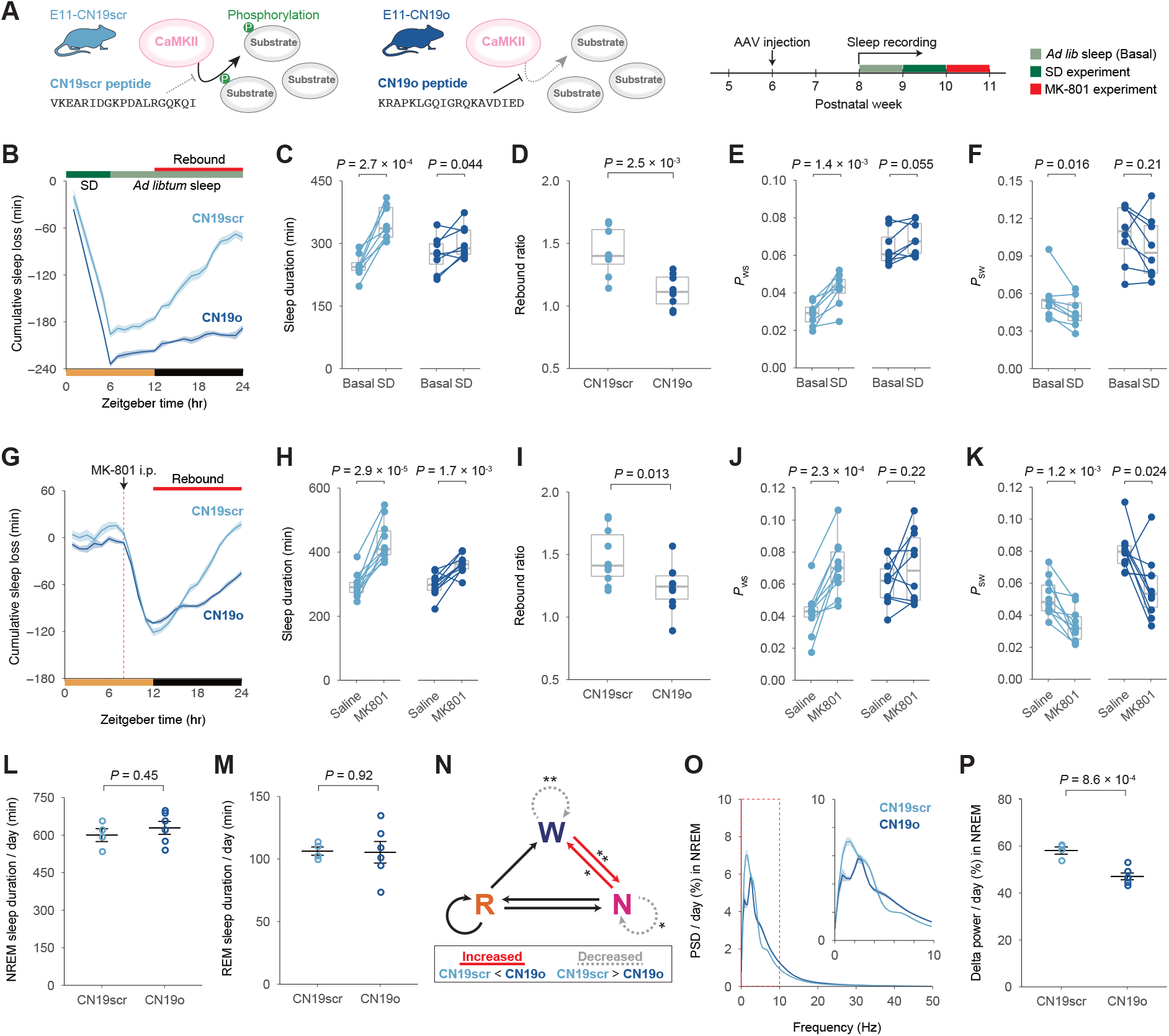
Inhibition of CaMKII activity in E11 neurons inhibits rebound sleep. (**A**) Diagram of CaMKII inhibition by CN19o peptide (left) and experimental schedule (right). (**B**) Cumulative sleep loss (SD – Basal) in E11-CN19o (navy) and E11-CN19scr (light blue) mice over 24 hours (*n* = 8 per group). Mice were allowed to behave freely for three days prior to SD, and the averaged value of sleep parameters is shown as “Basal.” (**C, E**, and **F**) Sleep duration, *P*_WS_, and *P*_SW_ on Basal and SD during ZT12–24. Student’s paired *t*-test was performed between Basal and SD within the groups. (**D**) Sleep rebound ratio (SD / Basal during ZT12-24). The groups were compared using Welch’s *t*-test. (**G**) Cumulative sleep loss (MK-801 – Saline) in E11-CN19o (navy) and E11-CN19scr (light blue) mice over 24 hours (*n* = 10 for each group). During the three days preceding the injections, the mice behaved normally. Each mouse received i.p. injection in turn: saline or 2 mg/kg MK-801 injection at ZT8, allowing at least 2 days between injections. (**H, J**, and **K**) Sleep duration, *P*_WS_, and *P*_SW_ on days with saline or MK-801 injection during ZT12–24. The Student’s paired *t*-test was used to compare the saline and MK-801 injections within the groups. (**I**) Sleep rebound ratio (MK-801 / Saline during ZT12–24). The groups were compared using Welch’s *t*-test. (**L** and **M**) Daily NREM sleep duration and REM sleep duration in E11-CN19scr (*n* = 4, light blue) and E11-CN19o (*n* = 6, navy) mice. The groups were compared using Welch’s *t*-test. (**N**) The differences in transition probabilities between wake, NREM, and REM sleep states in E11-CN19o mice versus E11-CN19scr mice. The solid red and dotted gray lines represent a significant increase and decrease, respectively. (**O**) Power spectral density (PSD) of the EEG during NREM sleep. The extracted diagram in the 0–10 Hz range is also shown in the top-right corner of the panel. (**P**) Normalized delta power (0.5–4 Hz) during NREM sleep. The groups were compared using Welch’s *t*-test. Error bar: SEM, \**P* < 0.05, \*\**P* < 0.01, \*\*\**P* < 0.001, n.s. = non significance.

We next performed a 6-hour SD on E11-CN19o and E11-CN19scr mice (**Fig. 3A**). Surprisingly, the E11-CN19o mice barely showed rebound sleep responses, whereas the E11-CN19scr mice clearly showed the responses following SD (**Fig. 3, B** to **F**; **fig. S8D**). After MK-801 administration, the E11-CN19o mice also showed lower rebound responses than the E11-CN19scr mice (**Fig. 3, G** to **K**; **fig. S8E**). These findings suggest that CaMKII kinase activity in E11 neurons is important for both basal sleep architecture and homeostatic sleep rebound.

We also recorded EEG and electromyogram (EMG) in E11-CN19o and E11-CN19scr mice. There was no significant difference in rapid eye movement (REM) sleep or non-REM (NREM) sleep duration between the E11-CN19o and E11-CN19scr mice (**Fig. 3, L** to **M**). The E11-CN19o mice, on the other hand, had higher transition probabilities between wake and NREM sleep states (i.e., increased W→N and N→W) and lower state stabilities (i.e., decreased W→W and N→N) (**Fig. 3N**). The level of delta (0.5–4.0 Hz) or slow oscillation (SO, 0.5–1.0 Hz) power during NREM sleep is correlated with the level of sleep need (*34, 35*). The delta power in E11-CN19o mice decreased significantly during NREM sleep (**Fig. 3, O** to **P**; **fig. S8, F** to **G**). The SO power did not change significantly but did tend to decrease (**Fig. 3O** and **fig. S8H**). These findings show that decreased CaMKII activity in the E11 neurons reduces NREM sleep quality (i.e., its stability and delta power).

### Elevation of CaMKIIα activity in E11 neurons induces rebound-sleep-like state

CaMKIIα is activated by Ca^2+^/calmodulin (CaM) binding and autophosphorylates threonine (T) 286 to maintain kinase activity even when Ca^2+^/CaM is not present (*36*) (**Fig. 4A**). Thus, the T286 phosphomimetic mutant (T286D), in which the threonine (T) is replaced with aspartic acid (D), becomes constitutively kinase-active form (*36*). To see if CaMKIIα activation in E11 neurons affects sleep phenotype, we expressed wild-type (WT) or T286D CaMKIIα in E11 neurons. Mice expressing WT CaMKIIα (E11-CaMKIIα (WT) mice) showed no differences in all sleep parameters when compared to E11-EGFP mice (**Fig. 4, B** to **D**). In contrast, E11-CaMKIIα (T286D) mice had significantly longer sleep duration, higher *P*_WS_, and lower *P*_SW_ than E11-EGFP and E11-CaMKIIα (WT) mice (**Fig. 4, B** to **D**). T286 non-phosphomimetic alanine (A) mutation (T286A) or kinase-dead mutation (K42R) adding to T286D (K42R:T286D) did not exhibit the changes of sleep parameters as observed in E11-CaMKIIα (T286D) mice (**fig. S9, A** to **C**). The expression of constitutively active (T287D) CaMKIIβ in E11 neurons also resulted in a rebound-sleep-like state as well as CaMKIIα activation (**fig. S9, D** to **F**).

**Fig. 4.**
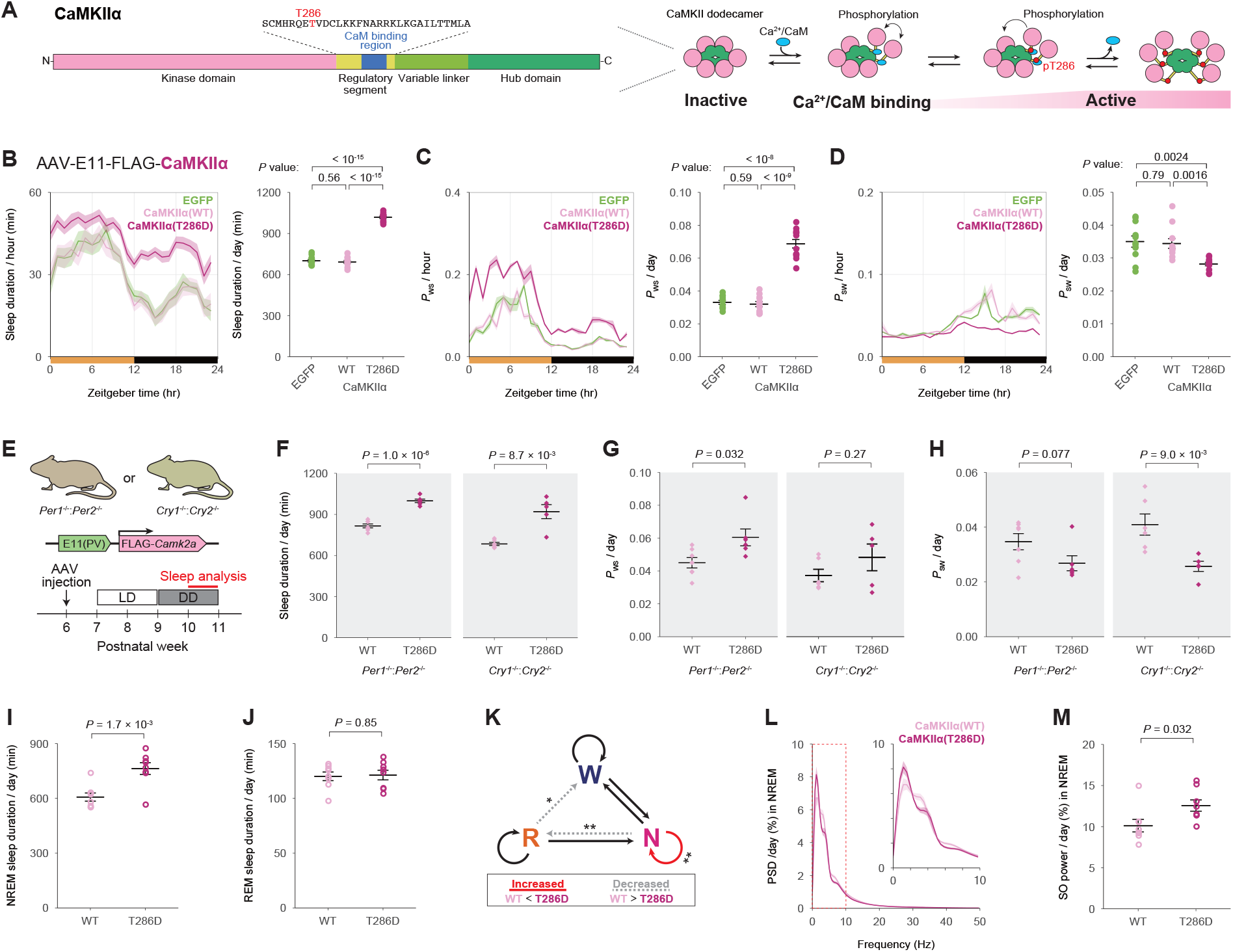
Elevation of CaMKIIα activity in E11 neurons induces rebound-sleep-like state. (**A**) Illustration of CaMKIIα protein sequences (left) and endogenous CaMKIIα activation mechanism (right). The phosphorylation of T286 residue in the regulatory segment is critical for CaMKIIα activity regulation. (**B, C**, and **D**) Daily sleep duration, *P*_WS_, and *P*_SW_ in E11-EGFP (*n* = 11, light green), E11-CaMKIIα (WT) (*n* = 11, pink), and E11-CaMKIIα (T286D) (*n* = 12, violet-red) mice. The groups were compared using Welch’s *t*-test. It should be noted that the significance level of *P*-values is corrected using the Bonferroni correction. (**E**) Diagram of CaMKII expression and experimental schedule in *Per* or *Cry* dKO mice under constant dark conditions. (**F, G**, and **H**) Daily sleep duration, *P*_WS_, and *P*_SW_ in the *Per* (left) or *Cry* (right) dKO mice expressing E11-CaMKIIα (WT) (*n* = 7, pink) or E11-CaMKIIα (T286D) (*n* = 6, violet-red). The groups were compared using Welch’s *t*-test. (**I** and **J**) Daily NREM sleep duration and REM sleep duration in E11-CaMKIIα (WT) (*n* = 8, pink) and E11-CaMKIIα (T286D) (*n* = 8, violet-red) mice. The groups were compared using Welch’s *t*-test. (**K**) The difference in transition probabilities between wake, NREM, and REM sleep states in E11-CaMKIIα (T286D) mice versus E11-CaMKIIα (WT) mice. The solid red and dotted gray lines represent a significant increase and decrease, respectively. (**L**) Power spectral density (PSD) of the EEG during NREM sleep. The extracted diagram in the 0–10 Hz range is also shown in the top-right corner of the panel. (**M**) Normalized slow-oscillation power (SO, 0.5–1Hz) during NREM sleep. The groups were compared using Welch’s *t*-test. Error bar: SEM, \**P* < 0.05, \*\**P* < 0.01, \*\*\**P* < 0.001, n.s. = non significance.

CaMKIIα (WT or T286D) was expressed in *Per1*^-/-^:*Per2*^-/-^ (*Per* dKO) or *Cry1*^-/-^:*Cry2*^-/-^ (*Cry* dKO) mice to see if the sleep-promoting effects of CaMKIIα activation in E11 neurons were dependent on the circadian clock genes (**Fig. 4E**). Under constant dark (DD) conditions, these dKO lines exhibit arrhythmic-circadian behavior (*22*). Regardless of circadian time, E11-CaMKIIα (T286D) mice in both dKO backgrounds kept under DD condition showed significantly longer sleep duration, higher *P*_WS_, and lower *P*_SW_ (**Fig. 4, F** to **H**; **fig. S9, G** to **L**).

Given the rebound-sleep-like phenotype observed in E11-CaMKIIα (T286D) mice (**Fig. 4, B** to **D**), it is reasonable to conclude that these mice have already acquired the saturated sleep need. To put this theory to the test, we performed a 6-hour SD on E11-CaMKIIα (T286D) mice. The E11-CaMKIIα (T286D) mice had rebound sleep characteristics even in basal sleep and only had marginal rebound sleep after the SD (**fig. S9, M** to **Q**). Similarly, after MK-801 administration, E11-CaMKIIα (T286D) mice showed lower rebound responses (**fig. S9, R** to **V**).

Furthermore, we recorded EEG and EMG in E11-CaMKIIα (WT) and E11-CaMKIIα (T286D) mice. The increased sleep duration in the E11-CaMKIIα (T286D) mice is due to an increase in NREM sleep (**Fig. 4I**), whereas REM sleep did not differ between groups (**Fig. 4J**). Consistent with this, E11-CaMKIIα (T286D) mice had higher NREM sleep stability (i.e., increased N→N) (**Fig. 4K**). The SO power was increased in E11-CaMKIIα (T286D) mice, but the delta power changes were not significant (**Fig. 4, L** to **M**; **fig. S10, A** to **C**). These findings show that constitutive CaMKIIα activation in E11 neurons increases both the quantity and quality of NREM sleep.

### CaMKII activity in E11 neurons encodes sleep need

CaMKII kinase activity in E11 neurons has similar effects on sleep homeostasis as E11-neuron activity manipulation. We thus hypothesized that CaMKII activation increases cortical PV-neuron activity, causing rebound sleep. To test this hypothesis, we performed double-immunostaining of PV and c-Fos proteins in brains of E11-CaMKIIα (T286D) mice. The isocortex contained more than three-quarters of the detected c-Fos+PV+ cells (**Fig. 5, A** to **B**). CaMKIIα (T286D) expression in E11 neurons selectively enhanced c-Fos expression in cortical PV neurons (**Fig. 5, C** to **E**) but had little effect on the overall number of c-Fos+ or PV+ cells (**fig. S11, A** to **B**). These findings imply that intracellular CaMKII activation increases PV-neuron activity.

**Fig. 5.**
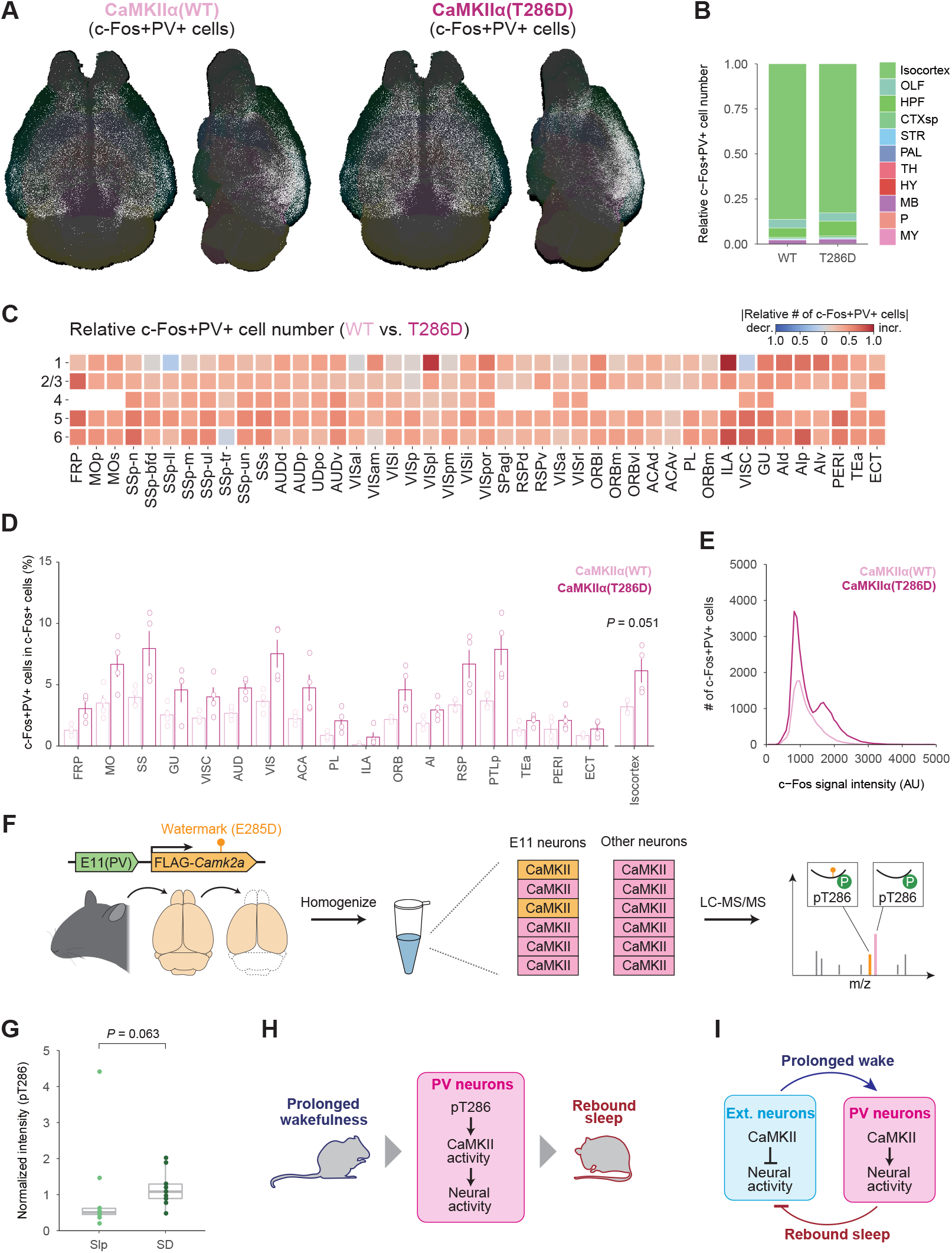
CaMKII activity in E11 neurons encodes sleep need. (**A**) Representative whole-brain views of dorsal and lateral double-positive (c-Fos+PV+) cell distribution (white). The brains of E11-CaMKIIα (WT) (left) and E11-CaMKIIα (T286D) (right) mice were collected. (**B**) Stacked percentage bar plots of c-Fos+PV+ cells in the E11-CaMKIIα (WT) and E11-CaMKIIα (T286D) brains (*n* = 4 for each group). The plot depicts the demographics of c-Fos+PV+ cells in major brain regions. (**C**) A region-wise heat map of relative cell number showing changes in c-Fos+PV+ cell number in the isocortex. The E11-CaMKIIα (T286D) brains were compared to the E11-CaMKIIα (WT) brains, with red indicating an increase and blue indicating a decrease. (**D**) The rate of double-positive (c-Fos+PV+) cells in c-Fos+ cells in each cortical region (left) or entire isocortex (right). The groups were compared using Welch’s *t*-test. (**E**) The cortical distribution of c-Fos signal intensity per c-Fos+PV+ cell in each group. (**F**) Experimental design for quantitative analysis of CaMKIIα T286 phosphorylation in E11 neurons. E11-CaMKIIα (T285D) mice were dissected, and homogenates were used for LC-MS/MS analysis. (**G**) Signal intensities of peptides derived from CaMKIIα (T285D) with phosphorylated T286 residues (*n* = 9 for each group). The intensities were normalized to the total CaMKIIα level, which was calculated using all quantifiable non-phosphorylated peptides derived from WT CaMKIIα. A two-sample Wilcoxon *t*-test was used to compare the groups. (**H**) CaMKII-dependent activity regulation of cortical PV neurons for homeostasis regulation. (**I**) Cortical feedback circuit model for sleep homeostasis regulation. Error bar: SEM. Brain region acronyms follow the ontology defined by the Allen Brain Atlas.

We next developed a system for measuring CaMKIIα T286 autophosphorylation using liquid chromatography-tandem mass spectrometry (LC-MS/MS) with watermark-mutant (E285D) CaMKIIα expression in E11 neurons (**Fig. 5F**), in which the glutamic acid (E) near T286 residue is replaced with aspartic acid (D). CaMKIIα (E285D) is distinguishable from all endogenous CaMKII isoforms (**fig. S12A**), allowing us to selectively quantify the T286 autophosphorylation level of CaMKIIα (E285D) expressed in the brain (**Fig. 5F**). We confirmed that CaMKIIα (E285D) expression and kinase activity were nearly identical to WT CaMKIIα (**fig. S12, B** to **C**). We also confirmed that no difference in any sleep parameters was observed between E11-CaMKIIα (WT) and E11-CaMKIIα (E285D) mice under basal and sleep-deprived conditions (**fig. S12, D** to **N**). We gave the E11-CaMKIIα (E285D) mice a 6-hour SD and collected their isocortex immediately afterward. Using the LC-MS/MS analysis, we revealed that T286 phosphorylation (pT286) level derived from CaMKIIα (E285D) increased in SD group compared to *ad libitum* sleep (Slp) group (**Fig. 5G**). These findings suggest that CaMKIIα kinase activity in cortical PV neurons reflects the level of sleep need.

## Discussion

We identified cortical PV neurons as a neural population responsible for homeostatic sleep rebound. Previous research has shown that subcortical and/or brain-stem regions regulate sleep homeostasis across the brain (*2*). However, as demonstrated by “local sleep,” brain-wide regulatory systems are unable to fully account for several aspects of sleep homeostasis (*3*). Recent research has revealed that cortical excitatory (pyramidal) neurons play a role in the regulation of sleep homeostasis (*37*). Our findings emphasize the importance of both cortical excitatory and inhibitory neurons in the regulation of sleep homeostasis. PV neurons can detect the synchronous firings of surrounding pyramidal neurons (*38*), and they can synchronize local circuit activity by inhibiting a number of surrounding pyramidal neurons (*8, 39*). Because synchronized cortical activity correlates with expected level of sleep need (*40*), cortical PV neurons may regulate the synchronicity of local cortical microcircuits and generate cortical slow waves in collaboration with other inhibitory neurons (*41*–*43*). The microcircuit is widespread: reciprocal connections between these two neural types are common across cortical areas/layers and species (*44*–*47*). The microcircuits may contribute to the local sleep homeostasis observed in mammalian cortex.

CaMKIIα/β-dependent phosphorylation responds well to sleep-wake states and sleep need, according to phosphoproteomic studies (*26*–*28*). Previous research has shown that genetic deletion of CaMKIIα/β or inhibition of CaMKII activity reduces daily sleep amount (*29, 30*). The kinase action in excitatory neurons promotes sleep, at least in part: expression of kinase-active CaMKIIβ mutant in excitatory neurons increases daily sleep amount (*30*). Our findings support the role of CaMKII in cortical PV neurons as a regulator of homeostatic sleep rebound. The molecular basis for the modulation is unknown, but it is possible that synaptic strength and/or intrinsic excitability in PV neurons are altered. Indeed, sleep-wake states influence electrophysiological activity and Ca^2+^ dynamics in cortical PV neurons (*20, 21*). CaMKIIα kinase activity is required for the sleep-promoting effects (**fig. S9, A** to **C**), but the downstream targets (i.e., substrates) are not the core circadian clock factors (**Fig. 4, F** to **H**; **fig. S9, G** to **L**). The voltage-dependent K^+^ channels K_v_3.1 and K_v_3.3 may be important candidates because double knockout of these channels changes the electrophysiological properties of PV neurons and impairs rebound sleep (*48*).

We propose that CaMKIIα is a key molecular factor in cortical PV neurons for encoding sleep need and inducing rebound sleep (**Fig. 7H**). The proposed mechanism may be responsible for local sleep homeostasis via the canonical negative feedback circuit in cortical inhibitory-excitatory neuron network (**Fig. 7I**). Such a regulatory motif for sleep homeostasis may be conserved across the animal kingdom: a distinct negative feedback circuit in flies is critical for sleep homeostasis regulation, with physiological properties modulated by sleep need (*49*). The design principle of the sleep homeostatic machinery may be shared by insects and mammals.

## Acknowledgments

We would like to express our gratitude to S. Tomita and A. Shimokawa for their assistance with the SSS and EEG/EMG recordings, S. Sato as well as K. Shimizu for assisting with AAV production, Y. Saito for assisting with brain sample preparation, M. Kuroda (International Research Center for Neurointelligence, the University of Tokyo) for LSFM imaging technical assistance, and T. Jin (RIKEN BDR) for supporting with NMR analysis. We would like to thank Enago (www.enago.jp) for the English language review.

## Author contributions

K.K., K.L.O., and H.R.U. designed the study; S. Shi, and E.A.S. helped with the study design; K.K. performed the EEG/EMG recording, SD experiments, chemogenetic experiments, most sleep measurements using the SSS system and analyzed the data; H.F. and D.T. performed part of the SSS measurements; K.K., K.L.O., D.T., and S.Y. designed and constructed the plasmids; K.K. and C.S. produced AAVs; K.K., T.M., C.S., and S. Shiono performed the brain sample preparation, LSFM imaging, and data analysis; K.K., K.L.O., and R.R.T. performed biochemical experiments and analyzed the data; Y.S. synthesized CNO for preliminary chemogenetic experiments; R.G.Y. established the improved FASTER methods for EEG/EMG sleep staging; D.T., J.Y.G., M.K., K.S., and H.K. established the ES-cell-derived mice; K.K. prepared the figures and draft manuscript; K.K., K.L.O., and H.R.U. wrote the manuscript with input from all co-authors.

## Data and materials availability

All data and materials used in this study are available from the corresponding author upon reasonable request.

## Supplementary Materials

Materials and Methods

figs. S1 to S12

References and Notes

