## Supplementary Materials for "Cortical parvalbumin neurons are responsible for homeostatic sleep rebound through CaMKII activation"

### Materials and Methods

#### *Mice*

The Animal Care and Use Committee of the Graduate School of Medicine in the University of Tokyo or the Institutional Animal Care and Use Committee of RIKEN BDR approved all animal care and experimental procedures. Mice were housed in humidity- and temperature-controlled conditions on a 12-hour light/dark (LD) cycle, with standard chow and water available *ad libitum* in groups, unless otherwise specified. This study used C57BL/6NJcl (WT) mice (CLEA Japan), *Per1*$^{-/-}$:*Per2*$^{-/-}$ (*Per* dKO) mice, and *Cry1*$^{-/-}$:*Cry2*$^{-/-}$ (*Cry* dKO) mice. *Per* dKO and *Cry* dKO mice were derived from ES cells using 3i methods and used in experiments without crossing (*50*, *51*). This study's mice were all male.

#### *Plasmids*

Except for CN19o expressing vectors, pAAV plasmids were built using pAAV-hSyn-DIO-hM3Dq-mCherry (Addgene #44361, kindly provided by Dr. Bryan Roth) (*52*). CN19o expressing vectors were constructed from another pAAV backbone kindly provided by Dr. Hirokazu Hirai (*30*). The promoter/enhancer, reporter/effector, WPRE, and polyA sequences were all present in all pAAV plasmids between ITRs (**fig. S5A**). The enhancers were placed upstream of a minimal human β-globin promoter (Addgene #83900, kindly provided by Dr. Gordon Fishell) (*53*). The Infusion HD Cloning Kit (Takara Bio) was used to clone these sequences.

The plasmids were then introduced into competent cells DH5α (TOYOBO) or NEB stable (New England Biolabs). The plasmids were purified using the Pure Yield Plasmid Midiprep System (Promega) and digested with *Xma*I (New England Biolabs) to ensure ITR integrity.

#### *Adeno-associated virus (AAV)*

In this study, we used an engineered AAV serotype, AAV-PHP.eB (*54*), and generated AAVs based on the previous reports (*55*). AAVpro 293T cells (Takara Bio) were transfected using Polyethylenimine (PEI, MW 25000, Polysciences) with three plasmids: pAAV (containing the gene(s) of interest), pUCmini-iCAP-PHP.eB (Addgene #103005, kindly provided by Dr. Viviana Gradinaru), and pHelper (Takara Bio). The medium from cultured AAVpro 293T cells was collected three days after transfection, and the cells and medium were harvested at least five days later. Centrifugation (2000 g, 15 min, and 25°C) was used to separate the harvested cells and medium. Freeze-thawing lysed the pellets, which were then treated with Benzonase Nuclease (Merck) or TurboNuclease (Accelagen) to digest non-packaged DNA. The lysates were mixed with 8% (wt/vol) polyethylene glycol (PEG MW 8000, MP Biomedical). AAVs were isolated from the salting-out supernatant using ultracentrifugation (58400 rpm, 2 hours 25 minutes, 18°C; Optima XE-90, Beckman Coulter; Type 70Ti Fixed-Angle Titanium Rotor, Beckman Coulter) with iodixanol (OptiPrep, Sigma). Before use, an AAV solution in Dulbecco's phosphate-buffered saline (DPBS) was concentrated using a centrifuge (AmiconUltra-15, Merck) and stored at 4°C.

The titration method was adapted from previous reports (*55*, *56*). Using quantitative PCR (qPCR), AAV titers were calculated as the number of effective vector genomes (vg) in AAV solution. To avoid contamination of non-packaged DNA, a portion of the AAV solution was treated with nuclease (50 U/ml, 1 hour, 37°C; Benzonase Nuclease or TurboNuclease). Proteinase K (30 µg/ml, 1 hour, 37°C; *N*-lauroylsarcosine in 3 mM NaCl, Fujifilm) was then added to the AAV solution to digest the viral capsid and release the viral genome. A general phenol/chloroform extraction and ethanol precipitation method was used to purify the viral genome. AAV titers (vg) were determined using qPCR with SYBR Green Master Mix (Thermo Fisher Scientific). The following primer sequences were used to amplify the target site (WPRE sequence):

WPRE_qPCR_fwd: 5′-CTGTTGGGCACTGACAATTC-3′

WPRE_qPCR_rev: 5′-GAAGGGACGTAGCAGAAGGA-3′

AAV vectors were administered to 6-week-old male mice via retro-orbital injection (**fig. S5A**). DPBS was used to adjust the injected AAV solution to the determined titer ($0.5–5.0 \times 10^{11}$ vg/mouse), and 100–150 µl/mouse was injected. Mice were anesthetized in ambient air with 1%–5% isoflurane prior to the injection. Mice were housed in their home cage for at least >2 weeks prior to behavioral experiments or brain sampling, unless otherwise noted, to allow time for gene expression.

***Respiration-based sleep measurements by Snappy Sleep Stager (SSS)***

SSS was used to perform sleep phenotyping based on respiration (*22*). The recordings were made in either WT male mice (3–12 weeks old) for the developmental analysis (details in below) or AAV-injected mice (8–10 weeks old) for other experiments. To measure respiratory flow, each experimental mouse was placed in the SSS chamber from its home cage. Unless otherwise specified, the recording condition was a 12-hour LD cycle with *ad libitum* food and water. Within two weeks, one set of recordings was completed. To avoid the first-night effect, data from the first day of the recording session were excluded from the analysis. The recordings of circadian KO mice were made under the DD condition for two weeks, and the data from the last six days was analyzed (**Fig. 4E**).

SSS data were divided into 8-second segments (epochs), with sleep/wake annotation performed after each epoch. The annotation rules and sleep parameters, such as sleep duration, $P_{WS}$, and $P_{SW}$, have previously been defined (*22*). $P_{WS}$ and $P_{SW}$ are the probabilities in an observed period from wake to sleep state (the tendency for sleep) and from sleep to wake state (the tendency for wakefulness). Because $N_{XY}$ denotes the number of transitions from X to Y (X, Y $\in$ {sleep (S), wake (W)}), the transition probabilities are independent of one another, and are calculated as follows:

$$P_{WS} = N_{WS} / (N_{WS} + N_{WW})$$

$$P_{SW} = N_{SW} / (N_{SW} + N_{SS})$$

$P_{WS}$ or $P_{SW}$ was treated as 1, when ($N_{WS} + N_{WW}$) or ($N_{SW} + N_{SS}$) was 0 (i.e., consecutive sleep or wake state in an observed period). The duration of a sleep or wake episode is calculated by taking the average of consecutive sleep or wake bouts, which is inversely related to $P_{SW}$ or $P_{WS}$, respectively.

Long-term recording with development lasted approximately 2 months in male mice, from P21 (just after weaning) to P84. During the recording, the SSS chambers were replaced with new ones every week, and sleep/wake annotation was performed on a weekly basis.

***Sleep measurements by EEG/EMG recording***

EEG and EMG surgery was performed 4–5 days after AAV injection (6–7 weeks old). A combination anesthetic containing 0.9 mg/kg medetomidine hydrochloride (Domitol, ZENOAQ), 4.8 mg/kg midazolam (Sandos, Novartis), and 6.0 mg/kg butorphanol (Vetorphale, Meiji Seika

Pharma) was used to anesthetize mice (*57*). Two EEG electrodes were placed in the skull over the cortex (1.5-mm right from midline, 1.5-mm anterior to the bregma; 1.5-mm right from the midline, 1.5-mm anterior to the lambda), and two EMG electrodes were placed in the neck muscle. Insulated wires connected the electrodes and were soldered to a pin header that was secured to the skull with dental cement. Mice were given 0.9 mg/kg atipamezole hydrochloride (Antisedan, ZENOAQ) after surgery and warmed to 37°C using a warming plate to promote anesthesia recovery. The mice were given at least ten days to recover from surgery.

Amplified EEG/EMG signals from 8-week-old male mice were recorded in the EEG/EMG chamber with food and water using Vital Recorder software (KISSEI COMTEC). For both EEG and EMG, the sampling frequency was 128 Hz. One recording session lasted less than a week. Data analysis began the day after data was collected. For further analysis, the EEG/EMG data recorded in KCD format was converted to EDF format.

For sleep recording, an in-house improved version of the FASTER method (*58*) was used to automatically annotate the stages (wakefulness, REM, and NREM sleep) from EEG/EMG data. EEG/EMG data were divided into 8-second segments (epochs), and annotation was performed after each epoch. Following FASTER's auto-staging, each epoch was manually corrected using the following criteria. The low-amplitude EEG and high amplitude EMG were used to determine wakefulness. NREM sleep was distinguished by high amplitude EEG with a low frequency wave (0.5–4 Hz) and a low EMG signal. The low-amplitude EEG with predominantly theta wave (6–10 Hz) and EMG atonia was used to stage REM sleep.

As previously reported (*59*), transition probabilities between wakefulness, NREM sleep, and REM sleep were calculated in the same way as $P_{WS}$ and $P_{SW}$ (see ***Respiration-based sleep measurements by SSS***). The transition from wakefulness to REM sleep ($P_{WR}$) was not depicted because it occurred so briefly.

For each epoch, the power spectral density (PSD) was calculated using fast Fourier transformation and Welch's averaging method (*60*). The spectral analysis excluded epochs that contained movement artifacts. Each PSD bin was calculated by averaging within each behavioral state and was expressed as a percentage of the total power over all frequencies. Delta or SO power was calculated during NREM sleep as percentages of delta (0.5–4 Hz) or SO (0.5–1 Hz) power over total EEG power per hour or day.

***Sleep measurements with SD***

SD was performed during ZT0–6 (the first half of light phase) in SSS recordings using an automated orbital shaker (TAITEC). Every mouse was moved from an SSS chamber to new cage on the shaker at ZT0 on the fourth day of recording. The shaker was programmed to turn on for 30 seconds every 80 seconds (30 second on, 50 second off) at around maximum speed of around 120 rpm. Following the SD session (at ZT6), each mouse was returned to the same SSS chamber as before, and the recording was resumed. The basal recording was made three days before the SD session.

***Sleep measurements with sleep pharmacological administration***

Natural saline was used to dissolve all chemicals used in pharmacological administration experiments (Otsuka pharma). Adjusting for individual mouse body weight, the single volume of injection was in the range of 150–250 µl. For MK-801 administration experiments, mice were given 0 or 2 mg/kg MK-801 maleate (Merck) i.p. injection at ZT8, with at least 1 day between injections.

For chemogenetic experiments, clozapine-*N*-oxide (CNO, Tocris) was used as the ligand of $G_q$-coupled DREADD hM3Dq and $G_i$-coupled DREADD hM4Di (*61*, *62*). For chemogenetic manipulation experiments under *ad libitum* sleep, mice expressing EGFP, hM3Dq, or hM4Di under the E11 enhancer were given i.p. injection of 0, 1, 3, or 5 mg/kg CNO at ZT12, with at least 2 days between injections. Mice expressing mCherry or hM4Di under the E11 enhancer were given i.p. injection of 0 or 0.25 mg/kg CNO at ZT12 for chemogenetic suppression experiments after SD using the above-mentioned method (see ***Sleep measurements with SD***).

***Whole-brain clearing, immunostaining, imaging, and analysis***

An improved version of the CUBIC-HV methods was used to clear and stain mouse brains (*24*). Mice were sedated with 100 mg/kg pentobarbital before being perfused transcardially with PBS and 4% paraformaldehyde (PFA). The brains were post-fixed in 4% PFA overnight at 4°C and stored in PBS until the clearing processes began. CUBIC-L (10% [wt/wt] N-butyldiethanolamine, 10% [wt/wt] Triton X-100 in water) was used to dilapidate the brains (*63*). Nuclear staining and immunostaining were performed after delipidation using nuclear staining dye (SYTOX Green, Thermo Fisher Scientific) and fluorescence-dye-conjugated antibodies. CUBIC-R+ (45% [wt/wt] antipyrine, 30% [wt/wt] Nicotinamide, 0.5% [vol/vol] N-butyldiethanolamine in water) was used to process brains for refractive index matching (*63*), followed by embedding in 1.5% agarose gel in CUBIC-R+.

All whole-brain imaging performed at International Research Center for Neurointelligence in the University of Tokyo. The microscope was outfitted with an Olympus 0.63X macro-zoom objective lens (Olympus) and a sCMOS camera (PCO). To achieve homogeneous resolution in XYZ axes, the microscope used an axially-swept light sheet mechanism.

Image analysis procedures on CUBIC-Cloud included cell detection, brain registration, and data quantification/visualization (*23*). The ilastik software was used to detect cells in 3D brain image data (*64*). Manual annotations were prepared from at least two brains with identical labeling conditions in some parts of the 3D images using ITK-SNAP software (*65*), and the classifier was trained using the manual annotation data. The XYZ position, mean fluorescence intensity, and volume of single cells at the whole brain level were obtained in CSV format by applying the trained classifier to the whole brain 3D image (*23*). The symmetric image normalization (SyN) algorithm was used to register the 3D brain image (*66*). Individual brains in CUBIC-Cloud were registered to CUBIC-Atlas brains using nuclear staining images and single cells displayed regional information (*67*). The CUBIC-Cloud platform (https://cubic-cloud.com/) was used to perform some data quantification and visualization.

***Whole-brain analysis of PV profile with development***

B6N WT male brains (P21, P28, P35, P42, P84, and P270; $n = 6$ for each age) were cleared, stained, imaged, and analyzed (see ***Whole-brain clearing, immunostaining, imaging, and analysis***). PV antibody (PV235, Swant; with anti-mouse IgG1 secondary Fab fragment conjugated with A594 [115-587-185, Jackson]) and SYTOX Green were used to stain the mouse brains.

***Whole-brain analysis of PV and c-Fos profile induced by sleep-wake perturbation***

B6N WT male mice (8 weeks old; $n = 6$ for Slp and SD groups) were individually housed in an SSS camber for 3 days prior to SD session for experiments. On the fourth day, mice in the SD group were sleep-deprived as described above (see ***Sleep measurements with sleep deprivation***), whereas mice in the Slp group were allowed to sleep *ad libitum* as much as they wanted. At ZT6 (just after the SD session), all mouse brains were sampled.

B6N WT male mice (8 weeks old; $n = 4$ for saline and MK-801 groups) were individually housed in an SSS camber for 3 days prior to i.p. administration of MK-801. On the fourth day, mice received i.p. injection of 0 or 2 mg/kg MK-801 maleate at ZT2. After the injection, mice were returned to the same SSS chamber for 1 day after the injection to monitor sleep phenotype. Mouse brains were collected for whole-brain analysis in a replicate experiment under identical conditions, except that mice were housed in the DD condition. The brains of mice were harvested two hours after i.p. injection (CT4).

To prepare brains for CaMKII activation in cortical PV neurons, the AAV.PHP.eB vector transducing E11-CaMKIIα (WT) or E11-CaMKIIα (T286D) was injected via retro-orbital sinus into B6N WT male mice (6 weeks old; $n = 4$ for each group). The mice were individually housed in an SSS camber for 1 week when they were 8 weeks old, to monitor their sleep phenotype. Following sleep phenotype, the brains of mice were collected.

In all PV and c-Fos analysis, collected mouse brains were cleared, stained, imaged, and analyzed as described above (see ***Whole-brain clearing, immunostaining, imaging, and analysis***). PV antibody (PV235, Swant; with anti-mouse IgG1 secondary Fab fragment conjugated with A594 [115-587-185, Jackson]), c-Fos antibody (2250S, CST; with anti-rabbit IgG secondary Fab fragment conjugated with Alexa 647 [111-607-008, Jackson]), and SYTOX Green were used to stain the mouse brains.

***In vitro assay for CaMKIIα expression and kinase activity***

HEK293T cells (ATCC) were transfected with pMU2-FLAG-*Camk2a* plasmids using PEI. After 24hours, the medium of cultured HEK293T cells was replaced with fresh culture medium. After 60 hours, the cells were lysed with 1 ml of cell lysis buffer (50 mM HEPES-NaOH pH 7.6, 150 mM NaCl, 0.5 mM $CaCl_2$, 1 mM $MgCl_2$, and 0.25% [v/v] NP-40) supplemented with 1% phosphatase inhibitor cocktail (Nacalai) and homogenized using the homogenizer (Hielscher). Cell lysates were collected and kept at −80°C.

Dot blot was used to estimate the relative expression levels of CaMKIIα in each cell lysate. Each cell lysate was diluted four times (1/4), then spotted on PVDF membrane (Merck). The membranes were reacted to anti-CaMKIIα antibody (SMC-124D, StressMarq) after blocking with Blocking One (Nacalai), followed by the reaction to the secondary antibody: anti-mouse IgG peroxidase (HRP)-conjugated antibody (W402B, Promega). HRP chemiluminescent reaction (Clarity Western ECL Substrate; ChemiDoc XRS+ system, Bio-Rad) was used to detect immunoreactivity of the blotted proteins. To ensure that the dot blot signals were within the linear range of detection, a cell lysate expressing the WT CaMKIIα was serially diluted (1, 1/2, 1/4, 1/8, 1/16, 1/32, 1/64, or 1/128) was spotted on the same membrane (data not shown).

The mobility shift assay was used to determine the CaMKII kinase activity in each cell lysate (LabChip EZ Reader II, PerkinElmer). Each diluted CaMKIIα cell lysate (1, 1/2, 1/4, 1/8, 1/16, 1/32, 1/64, or 1/128) was mixed with 0.25 mM ATP and 3.7 µM Fluorescently-labeled peptide substrate 11 (PerkinElmer) in the presence or absence of 0.50 µM CaM (Merck). The mixture was incubated at 37°C for 10 minutes, then at 98°C for 10 minutes to stop the reaction. CaMKII kinase activity is calculated by dividing the phosphorylated peptide signal by the total substrate peptide signal.

***Liquid chromatography-mass spectrometry/mass spectrometry (LC-MS/MS)***

B6N WT male mice (6 weeks old) were injected with the AAV.PHP.eB vector transducing E11-CaMKIIα (T285D) via retro-orbital sinus. The E11-CaMKIIα (T285D) mice were individually housed in an SSS camber for 1 week after they were 8 weeks old to monitor their sleep phenotype. The mice in the SD group were then sleep-deprived using the method described above (see ***Sleep measurements with SD***), whereas mice in the Slp group were allowed to sleep *ad libitum* as much as they wanted. The mice were killed by cervical dislocation at ZT6 (immediately following the SD session), and their cortical regions were quickly dissected and frozen in liquid nitrogen ($n$ = 9 for each group). The brain tissues were stored at −80°C to preserve phosphorylation state.

The PTS method was used to prepare samples for LC-MS/MS analysis (*30*, *68*). The frozen sample crusher (Tokken) was used to powder the brain tissues, and 5–10 mg of brain powder was homogenized in 0.5 ml of Solution B buffer (12 mM Sodium deoxycholate, 12 mM *N*-lauroylsarcosine sodium salt, 50 mM Ammonium hydrogen carbonate) supplemented with 1% phosphatase inhibitor cocktail (Nacalai), preheated at 98°C. Following a 15-minute incubation at 98°C, the samples were reduced with 9 mM dithiothreitol (Fujifilm) at room temperature for 30 minutes before being alkylated with 45 mM iodoacetamide (Merck) at room temperature for 30 minutes. The samples were then digested overnight at 37°C with 1 µg/ml of lysyl endopeptidase (Lys-C) (Fujifilm) and for 3–6 hours with 0.8 µg/ml of trypsin (Roche). Following digestion, the sample was acidified with 0.5% TFA and an equal volume of ethyl acetate was added. The sample was thoroughly mixed before being centrifuged (4000 rpm, 30 minutes, 25°C) to separate the peptide solution. Using a SpeedVac (Thermo Fisher Scientific), the peptide solution (an aqueous layer) was collected and dried.

The dried peptides were dissolved in a solution of 2% acetonitrile and 0.1% TFA. Each peptide solution was divided into two groups: one for individual samples, and the other for a mixture of each peptide solution as an internal control. Using a Sep-Pak C18 cartridge (Waters), the individual samples and internal control were trapped and desalted. The peptides on the cartridge were then dimethyl-labeled as previously described (*69*), with formaldehyde ($CH_2O$; Nacalai) added to the internal control (light label) or isotope-labeled formaldehyde ($CD_2O$; Cambridge Isotope Laboratories) added to the individual samples (medium label) together with $NaBH_3CN$ (Merck). The dimethyl-labeled peptides were eluted with a solution of 80% acetonitrile and 0.1% TFA. After that, an equal amount of light-labeled internal control was added to each medium-labeled individual sample, allowing us to compare the relative amount of peptides in the individual samples using internal control as a standard.

A portion of the mixture (10%) was subjected to LC-MS/MS analysis to determine the amount of CaMKIIα and total proteins. The remaining mixture (90%) was used to enrich the phosphorylated peptides using the High-Select Fe-NTA Phosphopeptide Enrichment Kit (Thermo Fisher Scientific) according to the manufacturer's protocol. All samples were SpeedVac dried before being dissolved in 2% acetonitrile and 0.1% TFA.

The samples were analyzed using an LC-MS system with Orbitrap Exploris 480 (Thermo Fisher Scientific) equipped with a nano HPLC system (Advance UHPLC; Bruker Daltonics) and an HTC-PAL autosampler (CTC Analytics) with a trap column (0.3 × 5 mm, L-column, ODS, Chemicals Evaluation and Research Institute). A gradient was used to separate samples using mobile phases

A (0.1% formic acid/$H_2O$) and B (0.1% formic acid and 100% acetonitrile) at a flow rate of 300 nl/ minute (4%–32% B for 190 minutes, 32%–95% B for 1 minute, 95% B for 2 minutes, 95%–4% B for 1 minute, and 4% B for 6 minutes) with a home-made capillary column (length of 200 mm and inner diameter of 100 µm) packed with 2-µm C18 resin (L-column2, Chemicals Evaluation and Research Institute). Several samples were separated using a different gradient (4%–36% B for 55 minutes, 36%–95% B for 1 minute, 95% B for 5 minutes, 95%–4% B for 1 minute, and 4% B for 8 minutes) without the phosphorylated peptide enrichment process. The eluted peptides were electrosprayed (2.1 kV) and subjected to positive ion mode analysis. Full MS spectra were obtained with a scan range of 350–1800 m/z and resolution of 60,000 FWHM at 200 m/z. MS2 spectra with 7,500 FWHM resolution at 200 m/z and a defined first mass at 120 m/z were obtained. Without the phosphorylated peptide enrichment process, samples were analyzed by data-dependent MS/MS acquisition, in which precursor ions (excluding isotopes of a cluster) above 5.0e3 intensity threshold with charge state ranging from 2+ to 7+ were selected for the MS/MS acquisition at 2.0 m/z isolation window during the 3 seconds interval between every full MS spectra acquisition. A dynamic exclusion of 20 seconds was used. Samples from the phosphorylated peptide enrichment process were analyzed by targeted MS/MS acquisition, where precursor ions matched with the dimethyl-labeled peptide, which included the phosphorylated T286 residue derived from the CaMKIIα and CaMKIIα (E285D) (i.e., light-WT 564.754 m/z, medium-WT 568.779 m/z, light-E285D 557.748, medium-E285D 561.773; z = 2+).

The raw data obtained were subjected to database searches (UniProt, reviewed mouse database as of October 5th, 2022) using the Sequest HT/Percolator algorithm running on Proteome Discoverer 2.5 (Thermo Fisher Scientific). Peptide cleavage was set to trypsin; missed cleavage sites were allowed to be up to two residues; peptide lengths were set to 6–144 amino acids; and mass tolerances were set to 40 ppm for precursor ions and 0.08 Da for fragment ions. Fixed modification included carbamidomethylation at cysteine and dimethylation [H (4) C (2), or 2H (4) C (2)] at lysine and the peptide N-terminus. Methionine oxidation was designated as a variable modification. A significance level of $q < 0.01$ was used. Precursor ion abundances for the phosphorylated T286 residue were manually checked and quantified using Qual browser function of the Xcalibur 4.4 software (Thermo Fisher Scientific). Proteome Discoverer 2.5 was used to calculate the abundances of the other precursor ions (Thermo Fisher Scientific).

***Statistical analysis***

Microsoft Excel for Mac version 16.49, R version 3.6.1., and Python version 3.8.2 were used for statistical analyses. For all tests, the significance level was set at < 0.05 (*$P < 0.05$, **$P < 0.01$, ***$P < 0.001$, n.s. = non-significance). The figure legends describe the statistical method and results for each comparison. The statistical analysis procedures were as follows.

To assess normality and homoscedasticity in two samples, the Shapiro test and *F*-test with a significance level of 0.05 were used. When two groups had normal distributions with equal variance, the Student's *t*-test was used for independent samples; and the Student's paired *t*-test was used for paired samples. When two groups had normal distributions with unequal variance, Welch's *t*-test was used; otherwise, two-sample Wilcoxon *t*-test was used.

To evaluate normality and homoscedasticity in more than three samples, the Kolmogorov–Smirnov test (KS-test) and Bartlett's test at the significance level of 0.05 were used. When all groups were normally distributed with equal variance, the Student's *t*-test with Bonferroni correction was used; when two groups had normal distributions but not equal variance, the Welch's *t*-test with Bonferroni correction was used; otherwise, the Steel test was used.

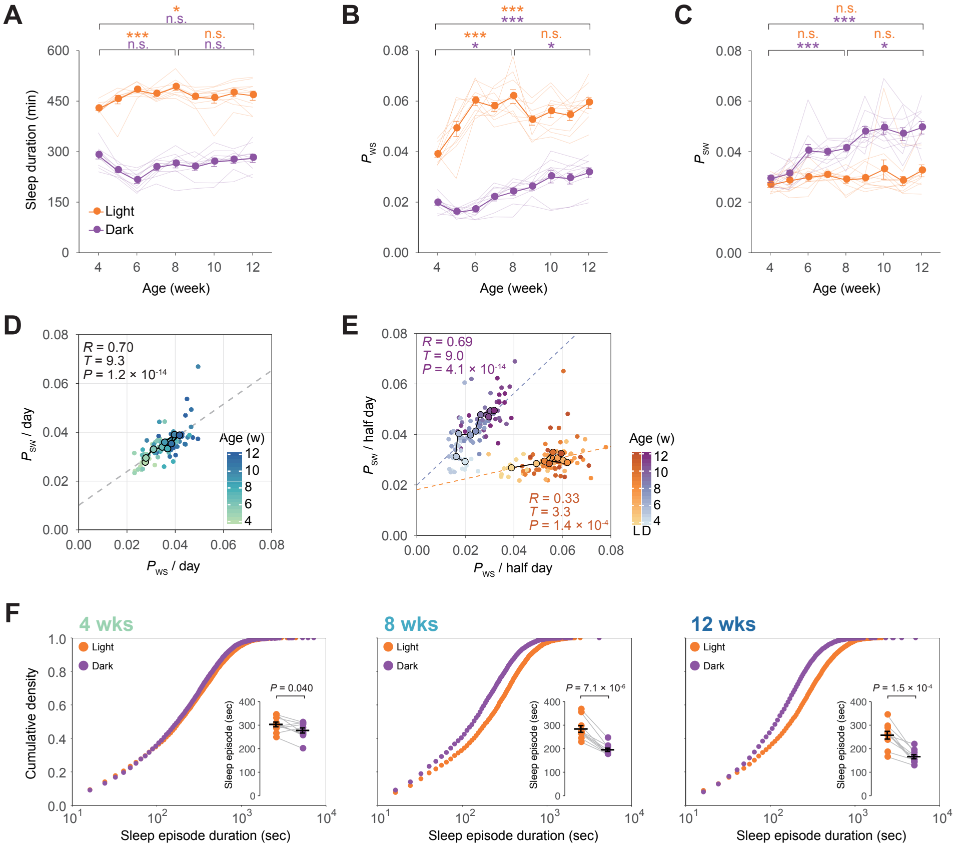


**fig. S1. Analysis of developmental changes in sleep architecture.**

(**A**, **B**, and **C**) Half-day sleep duration, $P_{\mathrm{ws}}$, and $P_{\mathrm{sw}}$ in the developing mice ($n$ = 10). At each age, the orange and purple points represent the mean sleep parameters during the light phase (first half of the day) and the dark phase (second half of the day). Each line (light orange and purple) represents the sleep parameters of a single developing mouse during the phases. Welch's $t$-test with Bonferroni correction was used on animals aged 4, 8, and 12 weeks. (**D** and **E**) Correlation diagram of $P_{\mathrm{WS}}$ and $P_{\mathrm{SW}}$ in developing mice over a day and half-day. The mean of $P_{\mathrm{WS}}$ and $P_{\mathrm{SW}}$ at each age is shown by black-fringed dots, while other dots show the $P_{\mathrm{WS}}$ and $P_{\mathrm{SW}}$ of individual developing mice, whose color corresponds to their age. At each age, orange and purple points represent the parameters during the light phase and the dark phase, respectively. The regression line is represented by each dashed line. The diagrams depict the correlation coefficient ($R$), $T$-, and $P$-value. (**F**) Cumulative frequency plot of sleep episodes. The ages of 4, 8, and 12 weeks are chosen representative data. Welch's $t$-test was used to compare the mean sleep episodes in light and dark phases at 4, 8, and 12 weeks old ($n$ = 10, at each age). Error bar: SEM, $^{*}P < 0.05$, $^{**}P < 0.01$, $^{***}P < 0.001$, n.s. = non significance.

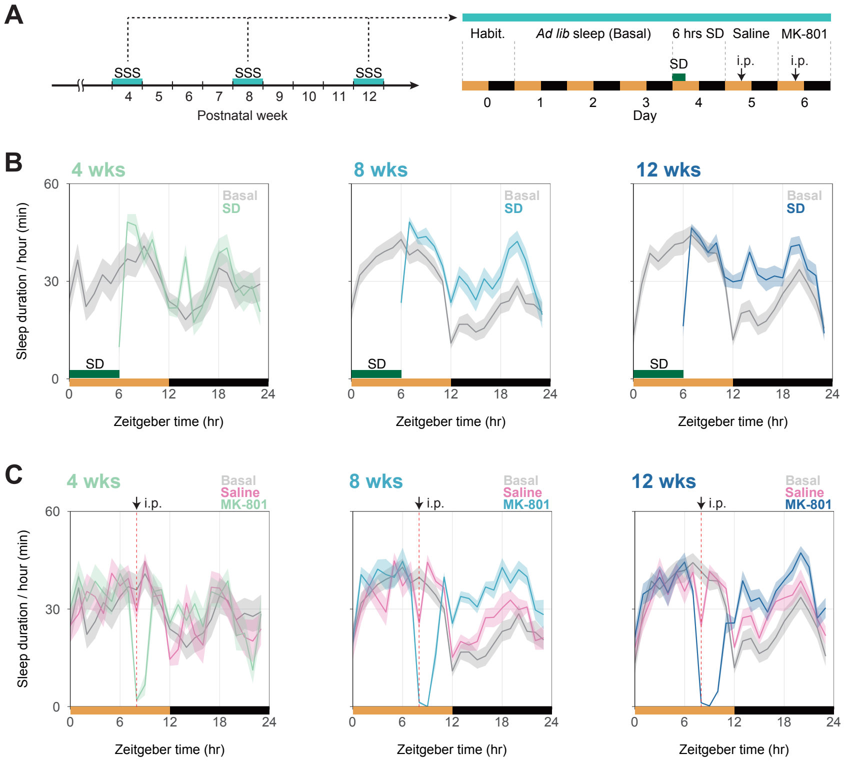


**fig. S2. Analysis of developmental changes in sleep homeostasis.**

(**A**) Schematic representation of sleep deprivation (SD) experiments performed on developing mice (*n* = 12) at 4, 8, and 12 weeks of age using the SSS system. Mice were freely behaving during the first 3 days, and the averaged value of sleep parameters is shown as "Basal." On the fourth day, SD was performed for 6 hours from ZT0 to ZT6. On the fifth (Saline) and sixth (MK-801) days, respectively, saline and 2 mg/kg MK-801 were administered at ZT8. (**B** and **C**) Hourly sleep duration over 24 hours in developing mice (n = 12) at 4, 8, and 12 weeks old during sleep deprivation and MK-801 administration. In the panels (B) and (C), the basal (gray line) is the same.

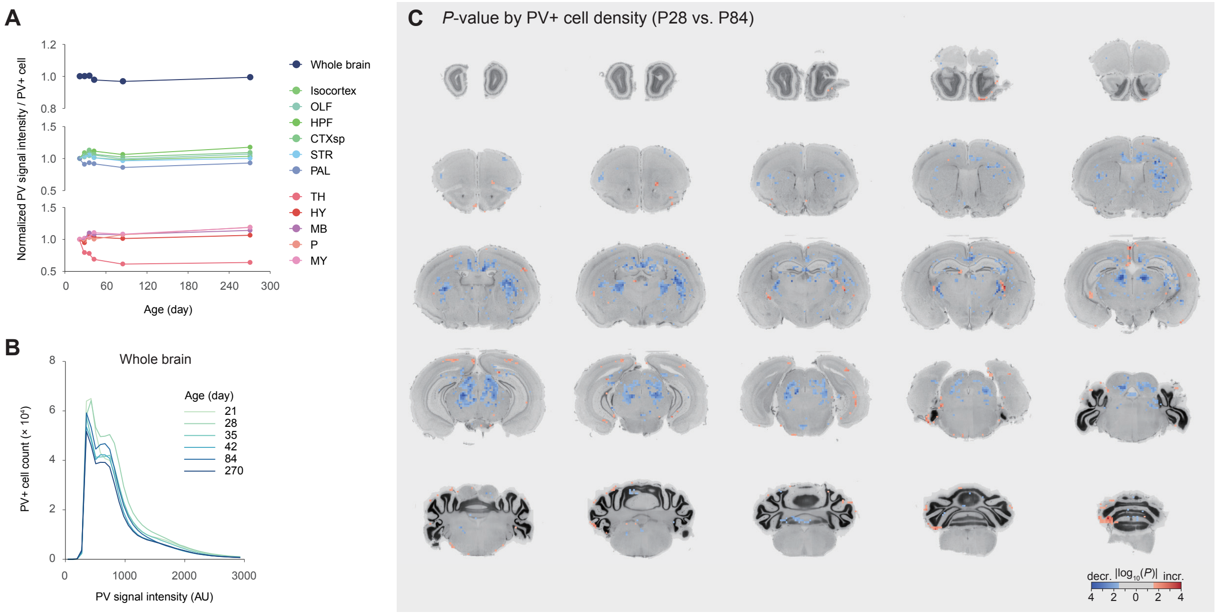


**fig. S3. Whole-brain analysis of PV neurons across post-weaning development.**

(**A**) The mean of PV signal intensity per PV+ cell in the whole brain (top), cerebral regions (middle), and brain-stem regions (bottom) across post-weaning development ($n$ = 6 for each age). Each point represents the mean value of each age normalized by that of P21. (**B**) The distribution of PV signal intensity per PV+ cell over the entire brain at each age. (**C**) A virtual cortical slice depicts a voxel-wise $P$-value heat map of PV+ cell density in comparison to P28 and P84 brains. P28 and P84 brains were tested using Welch's $t$-test. Red and blue colors represent a significant increase and decrease in P84 brains compared to P28 brains, respectively. Gray was assigned to regions with no significance ($P > 0.05$). (**D**) The mean of PV+ cell density in selected subcortical and brain-stem regions during post-weaning development ($n$ = 6 for each age). Each point represents the mean value of each age normalized by that of P21, and the colors displayed correspond to the brain regions depicted in panel (A).

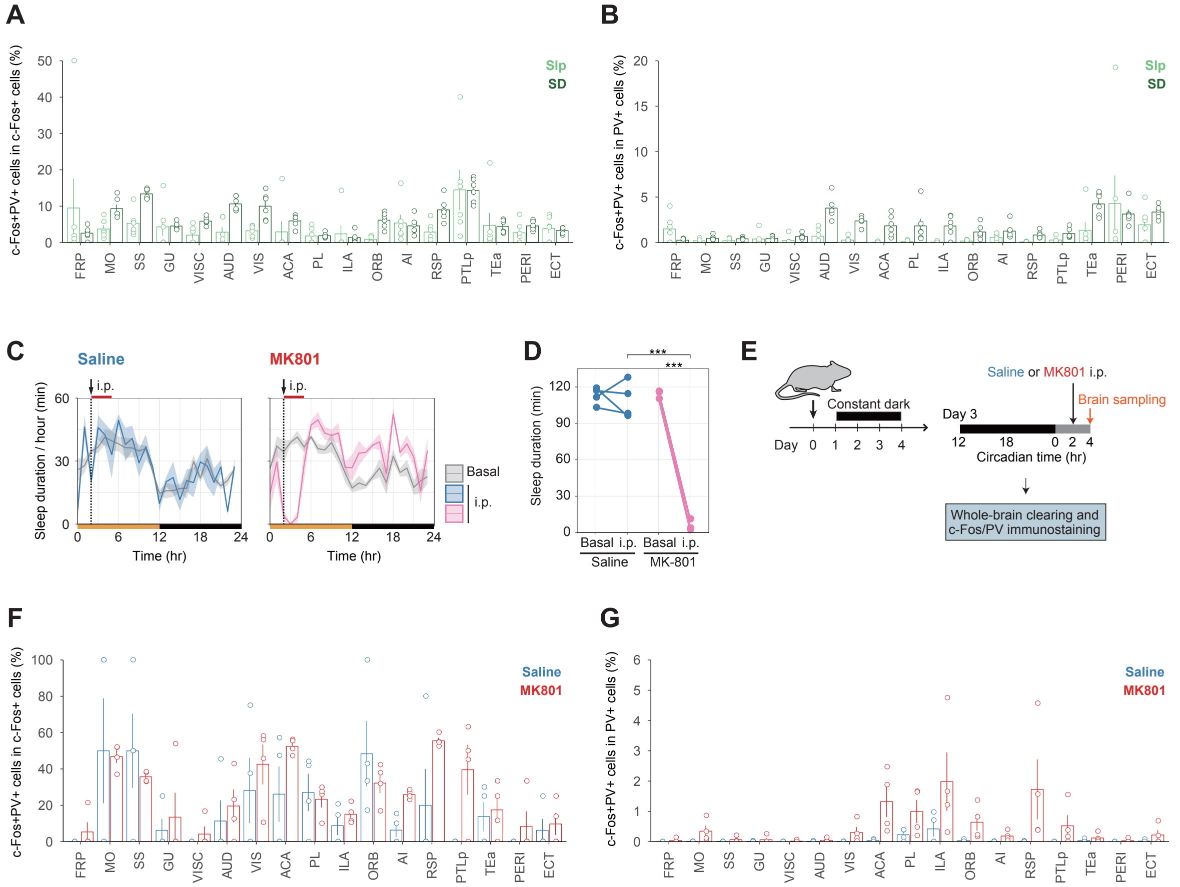

**fig. S4. Analysis of cortical PV-neuron activity upon sleep deprivation or MK-801 administration.**

(**A** and **B**) Rate of double-positive (c-Fos+PV+) cells in c-Fos+ cells or PV+ cells in each region of the isocortex after SD. (**C**) Hourly sleep duration over 24 hours in MK-801 administration experiment (*n* = 4 for each group). Mice were freely behaving during the first 3 days, and the averaged value of sleep parameters is shown as "Basal." On the fourth day, mice were given i.p. injection of saline or 2 mg/kg MK-801 at ZT2. (**D**) Sleep duration during ZT2–5 on both the Basal and the injection days. Within or between groups, Welch's *t*-test with Bonferroni correction was used. (**E**) Experiment design for whole-brain analysis of c-Fos+ and/or PV+ cell distribution in response to MK-801 administration. Individual mice were housed in constant darkness (DD) from three days prior to i.p. injection. All brains of control (Saline) and MK-801-administered (MK-801) mice were collected at CT4 (*n* = 4 for each group) after the injection at CT2. Each set of whole-brain images was uploaded to the CUBIC-Cloud and registered to the CUBIC-Atlas brain. (**F** and **G**) Rate of double-positive (c-Fos+PV+) cells in c-Fos+ cells or PV+ cells in each region of the isocotex after MK-801 administration. Error bar: SEM, $*P < 0.05$, $**P < 0.01$, $***P < 0.001$, n.s. = non significance. The Allen Brain Atlas ontology is used to define brain region acronyms.

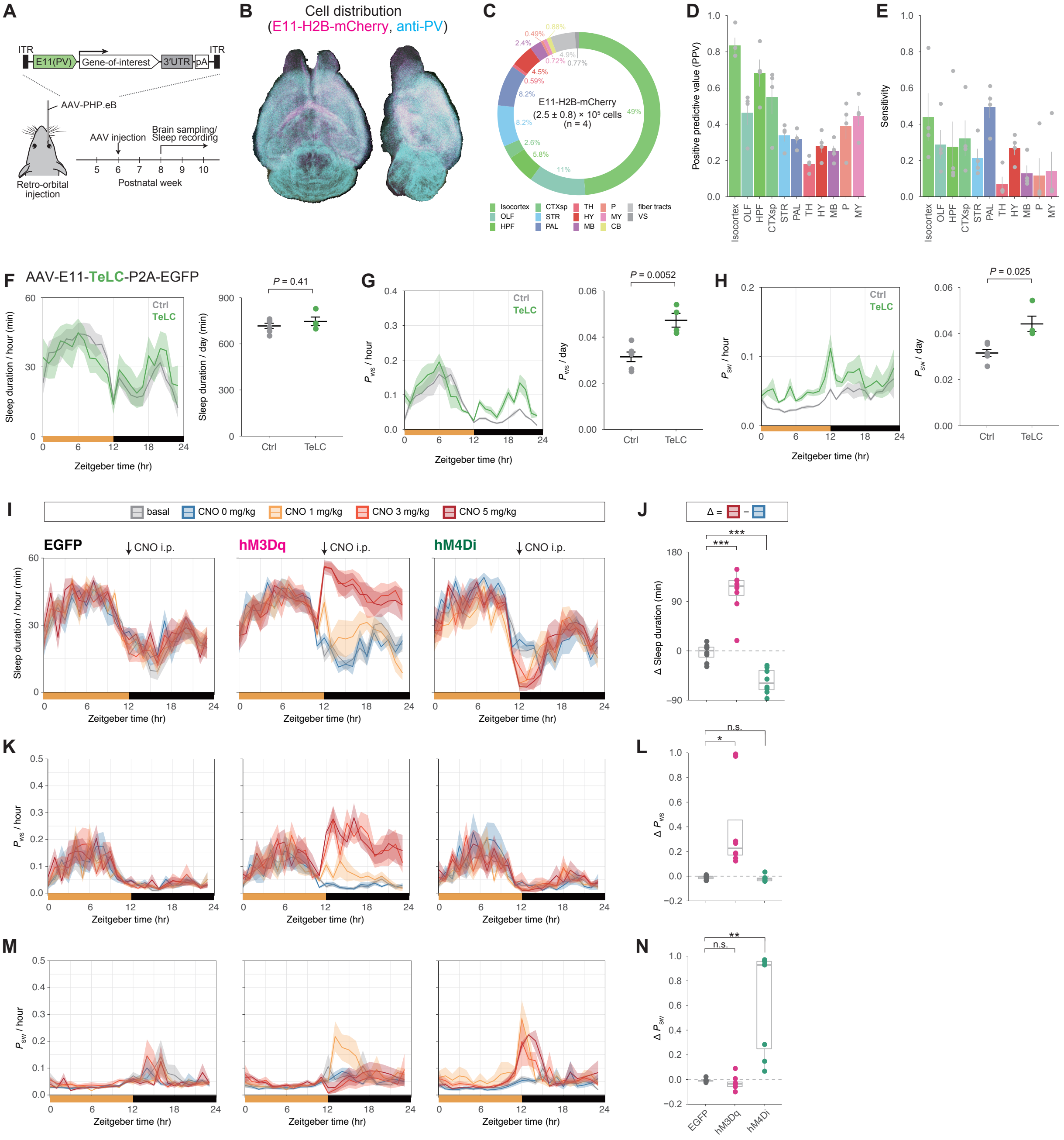

**fig. S5. Chronic or acute manipulation of E11-neuron activity changes the sleep-wake states.**
(**A**) Experimental design for AAV-based gene expression under E11 enhancer. To allow time for gene expression, AAV-injected mice were housed in their home cage for > 2 weeks before behavioral experiments or brain sampling. (**B**) Representative whole-brain views of dorsal and lateral E11-H2B-mCherry (magenta) and anti-PV (cyan) signals. (**C**) A donut plot depicting the whole-brain distribution of E11-H2B-mCherry signals ($n$ = 4). The plot depicts the demographics of mCherry-positive (mCherry+) cells in major brain regions. (**D**) The positive predictive value was calculated by dividing the number of colocalized (mCherry+PV+) cells in the major brain regions by the total number of mCherry+ cells in the brain. (**E**) Sensitivity was calculated by

dividing the number of colocalized (mCherry+PV+) cells by the total number of PV+ cells in the major brain regions. (**F**, **G**, and **H**) Daily sleep duration, $P_{WS}$, and $P_{SW}$ in Ctrl (*n* = 6) and E11-TeLC (*n* = 4) mice. The groups were compared using Welch's *t*-test. (**I**, **K**, and **M**) Hourly sleep duration, $P_{WS}$, and $P_{SW}$ over 24 hours on the basal day (gray; the mean of two days with no injection) and days with CNO injection. Each mouse received i.p. injection in turn: 0 (blue), 1 (gold), 3 (red), or 5 (dark red) mg/kg CNO injection at ZT12 with at least 2 days between injections (*n* = 8 for each group). (**J**, **L**, and **N**) The differences (Δ) in sleep duration, $P_{WS}$, and $P_{SW}$ between the 0 and 5 mg/kg CNO conditions during ZT12–15. Each dot represents a single mouse. Welch's *t*-test with Bonferroni correction was used to compare the groups to the E11-EGFP mouse group. Error bar: SEM, $*P < 0.05$, $**P < 0.01$, $***P < 0.001$, n.s. = non significance. Brain region acronyms follow the ontology defined by the Allen Brain Atlas.

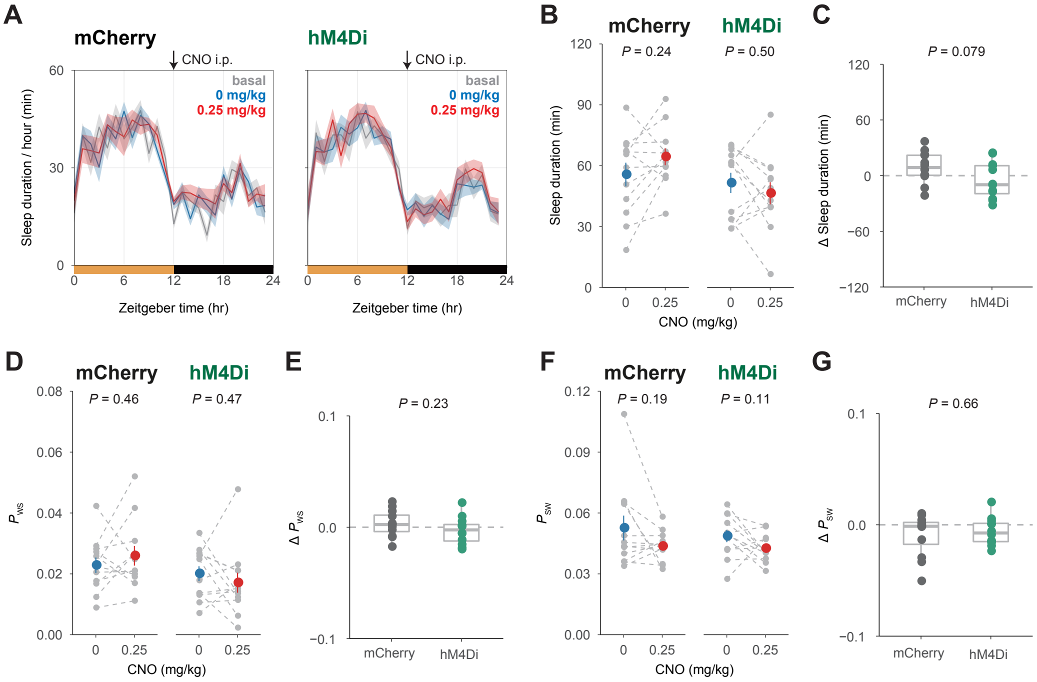


**fig. S6. Suppression of E11 neurons by the low-dose CNO administration barely affects basal sleep architecture.**

(**A**) Hourly sleep duration over 24 hours on the baseline day (gray; the mean of two days with no injection) and days with CNO injection. Each mouse received i.p. injection in turn: 0 (blue) or 0.25 (red) mg/kg CNO injection at ZT12, with at least 2 days between injections (*n* = 12 for each group). (**B**, **D**, and **F**) Sleep duration, $P_{WS}$, and $P_{SW}$ on the days with CNO injection during ZT12–15. Each gray dot represents an individual mouse, while the colored dots represent the mean in each condition. Individual mouse's dots are connected by gray-dashed lines. Welch's *t*-test was used to compare conditions within groups. (**C**, **E**, and **G**) The differences (Δ) in sleep duration, $P_{WS}$, and $P_{SW}$ between days with 0 and 0.25 mg/kg CNO injections during ZT12–15. The group were compared using Welch's *t*-test. Error bar: SEM.

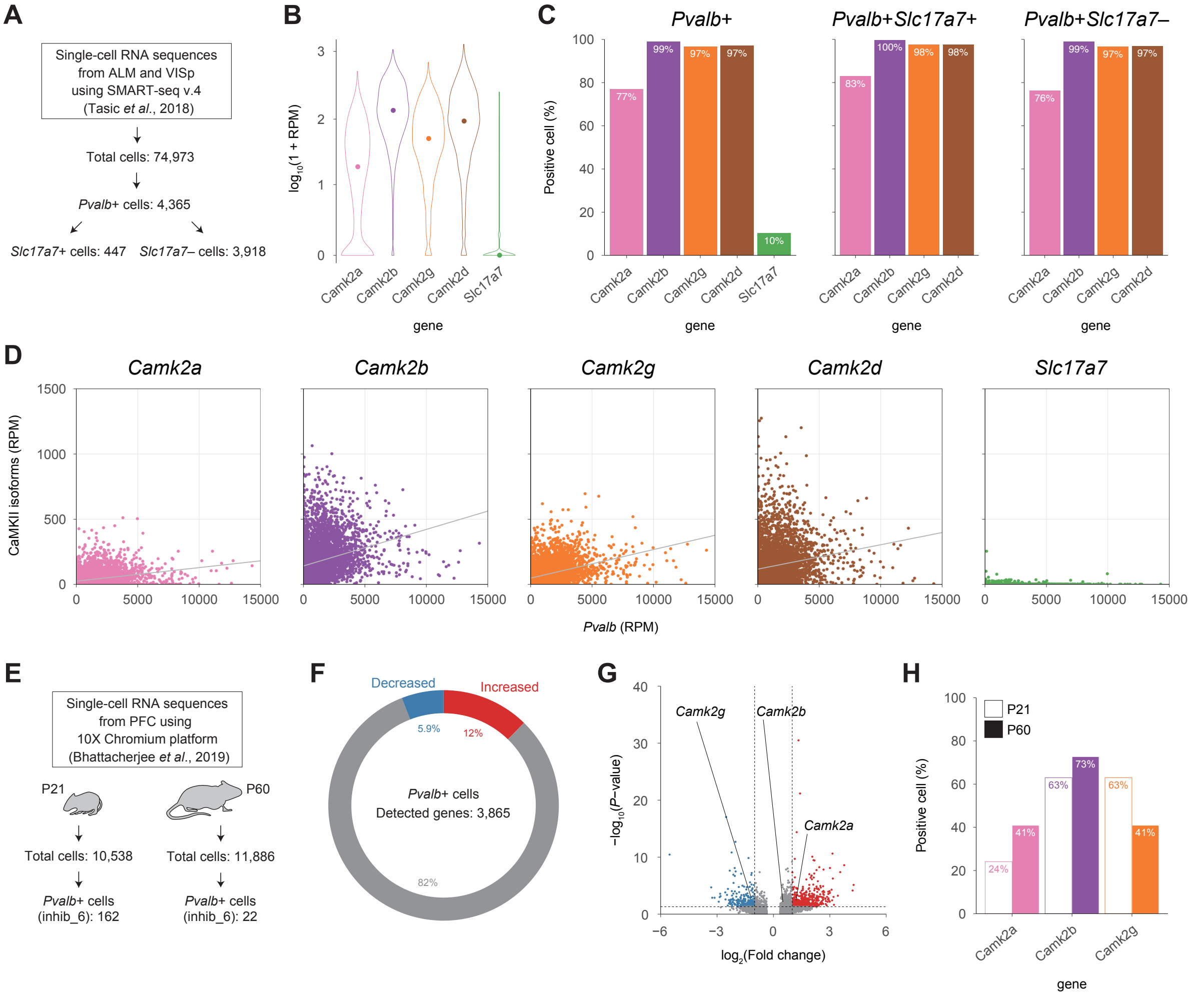


**fig. S7. Single-cell RNA sequence analysis of CaMKII expression in cortical *Pvalb*+ cells.**
(**A**) Single-cell RNA sequence analysis in the adult (P53–59) mouse cortex (ALM or VISp) using the Allen Brain Institute data (*31*). *Pvalb*+ cells were the cells in the data that were assigned the *Pvalb* subclass. *Pvalb*+ cells were further subdivided into *Slc17a7*-positive (*Pvalb*+*Slc17a7*+) and negative (*Pvalb*+*Slc17a7*−) cells (*Slc17a7*: glutamatergic neuron marker gene). (**B**) Violin plots depict gene expression of all CaMKII isoforms and *Slc17a7* in *Pvalb*+ cells ($n = 4{,}365$). On a $\log_{10}$ scale, expression values (reads per million, RPM) are displayed. The medians are indicated by colored dots. (**C**) Positive cell (Reads > 0) rate in *Pvalb*+ ($n$ = 4,365, left), *Pvalb*+*Slc17a7*+ ($n$ = 447, center), and *Pvalb*+*Slc17a7*− ($n$ = 3,918, right) cells. (**D**) Correlation diagram of the *Pvalb* and each selected gene (CaMKII isoforms and *Slc17a7*) expression values (RPM) in *Pvalb*+ cells ($n$ = 4,365). Each dot represents a single cell. The gray lines represent the regression lines. (**E**) Method for comparing single-cell RNA sequences in the adolescent (P21) and young-adult (P60) mouse prefrontal cortex (PFC) using data from the Yi Zhang group (*32*). The cells in the data assigned to the Inhib-6 cluster were dubbed *Pvalb*+ cells. (**F** and **G**) Donut plot and volcano plot of up-regulated (red) or down-regulated (blue) gene expression ($n$ = 3,865 genes) in *Pvalb*+ cells from P21 ($n$ = 162 cells) to P60 ($n$ = 22 cells) mouse PFC (cutoff: averaged fold change > 2 and *P*-value < 0.05). The volcano plot highlights CaMKII isoform genes that have been identified. (**H**)

Positive cell (Reads > 0) rate of each CaMKII isoform gene in *Pvalb*+ cells of mouse PFC at P21 ($n$ = 162 cells) and P60 ($n$ = 22 cells). P21 and P60 are represented by edged and filled bars, respectively. Abbreviations: ALM, the anterior lateral motor cortex; VISp, the primary visual cortex.

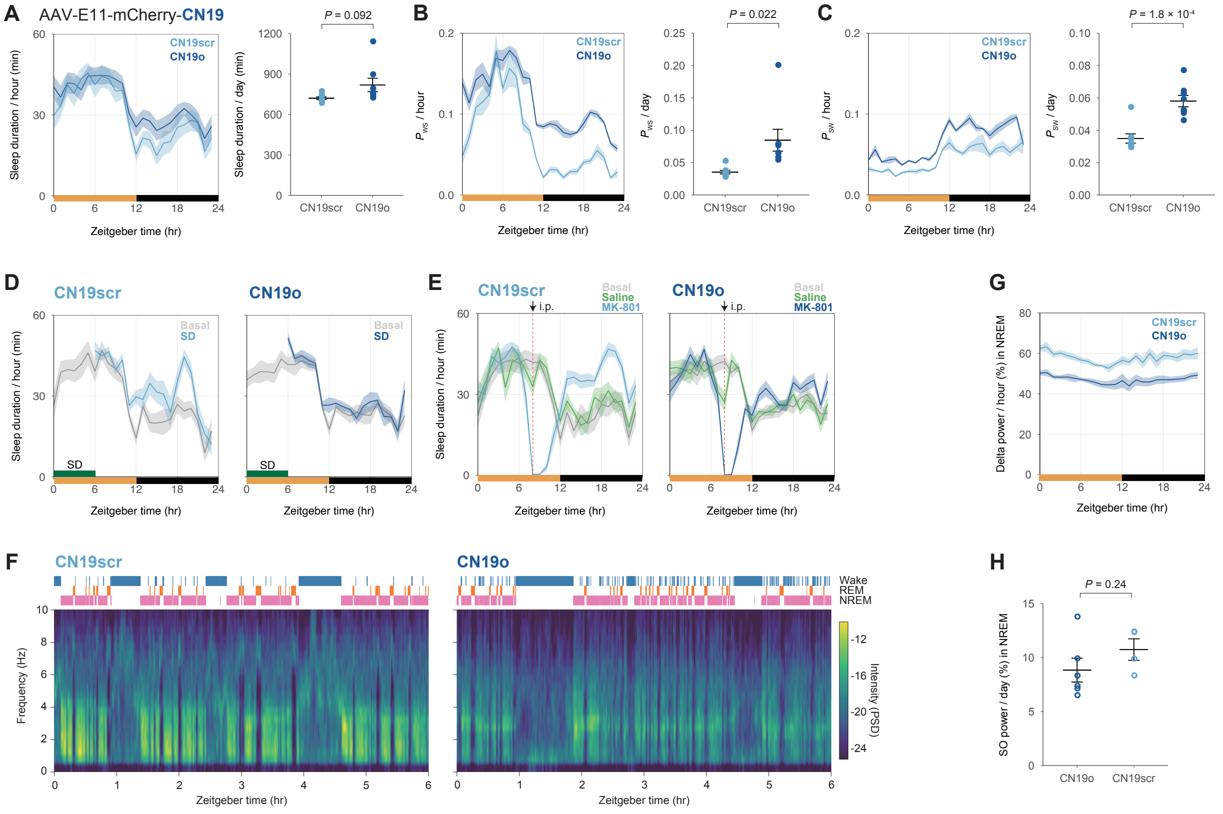


**fig. S8. Inhibition of CaMKII kinase activity in E11 neurons affects sleep quality not quantity.** (**A**, **B**, and **C**) Daily sleep duration, $P_{WS}$, and $P_{SW}$ in E11-CN19o (navy) and E11-CN19scr (light blue) mice ($n$ = 8 for each group). The groups were compared using Welch's $t$-test. (**D** and **E**) Hourly sleep duration over 24 hours in SD and MK-801 administration experiments. Each mouse received i.p. injection in turn: saline or 2 mg/kg MK-801 injection at ZT8, with at least 2 days between injections. Mice were allowed to behave freely for three 3 days prior to SD or the first injection, and the averaged value of sleep parameters is shown as "Basal." (**F**) Representative sleep-wake features of E11-CN19scr (left) and E11-CN19o (right) mice are shown as hypnograms (top) and heat maps of EEG spectra (bottom). (**G**) Hourly normalized delta power (0.5–4 Hz) during NREM sleep in E11-CN19scr ($n$ = 4, light blue) and E11-CN19o ($n$ = 6, navy) mice. (**H**) Normalized slow-oscillation (SO) power (0.5–1 Hz) during NREM sleep. The groups were compared using Welch's $t$-test. Error bar: SEM.

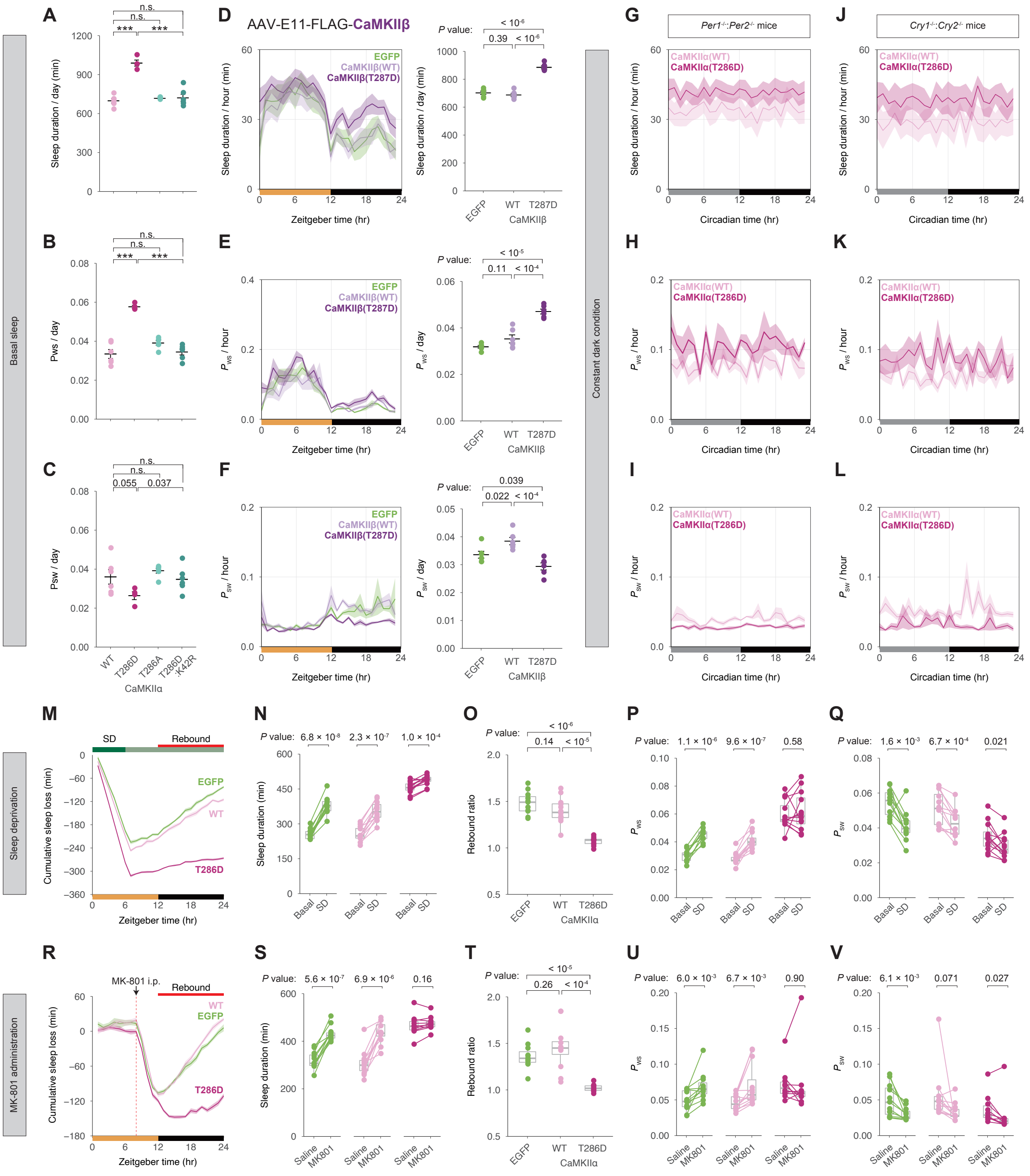


**fig. S9. Elevation of CaMKIIα or β kinase activity in E11 neurons increases daily sleep amount and consolidates sleep state.**

(**A**, **B**, and **C**) Daily sleep duration, $P_{\mathrm{WS}}$, and $P_{\mathrm{SW}}$ in E11-CaMKIIα mutant mice ($n$ = 6 for each group, except for T286D ($n$ = 4)). Welch's $t$-test with Bonferroni correction was used to compare the groups to E11-CaMKIIα (WT) or E11-CaMKIIα (T286D) mice. (**D**, **E**, and **F**) Daily sleep duration, $P_{\mathrm{WS}}$, and $P_{\mathrm{SW}}$ in E11-EGFP (light green), E11-CaMKIIβ (WT) (light purple), and E11-CaMKIIβ (T287D) (purple) mice ($n$ = 6 for each group). The groups were compared using Welch's *t*-test. It should be noted that the significance level of *P*-values is corrected using the Bonferroni correction. (**G**, **H**, and **I**) Hourly sleep duration, $P_{WS}$, and $P_{SW}$ over 24 hours in the *Per* dKO mice expressing E11-CaMKIIα (WT) (*n* = 7, pink) or E11-CaMKIIα (T286D) (*n* = 6, violet-red). (**J**, **K**, and **L**) Hourly sleep duration, $P_{WS}$, and $P_{SW}$ over 24 hours in the *Cry* dKO mice expressing E11-CaMKIIα (WT) (*n* = 6, pink) or E11-CaMKIIα (T286D) (*n* = 5, violet-red). (**M**) Cumulative sleep loss (SD – Basal) in E11-EGFP (*n* = 11, light green), E11-CaMKIIα (WT) (*n* = 12, pink), and E11-CaMKIIα (T286D) (*n* = 12, violet-red) mice over 24 hours. Mice were allowed to behave freely for three days prior to SD, and the averaged value of sleep parameters is shown as "Basal." (**N**, **P**, and **Q**) Sleep duration, $P_{WS}$, and $P_{SW}$ on Basal and SD during ZT12–24. Within the groups, Student's paired *t*-test was used to compare Basal and SD. It should be noted that the significance level of *P*-values is corrected using the Bonferroni correction. (**O**) Sleep rebound ratio (SD / Basal during ZT12–24). The groups were compared using Welch's *t*-test. It should be noted that the significance level of *P*-values is corrected using the Bonferroni correction. (**R**) Cumulative sleep loss (MK-801 – Saline) in E11-EGFP (*n* = 11, light green), E11-CaMKIIα (WT) (*n* = 12, pink), and E11-CaMKIIα (T286D) (*n* = 10, violet-red) mice over 24 hours. The mice were allowed to behave freely for three days prior to the first injection, and the averaged value of sleep parameters is shown as "Basal." Each mouse received i.p. injection in turn: saline (blue) or 2 mg/kg MK-801 injection at ZT8, allowing at least 2 days between injections. (**S**, **U**, and **V**) Sleep duration, $P_{WS}$, and $P_{SW}$ on the days with saline or MK-801 injection during ZT12–24. Student's paired *t*-test was used to compare the saline and MK-801 injections within the groups. It should be noted that the significance level of *P*-values is corrected using the Bonferroni correction. (**T**) Sleep rebound ratio (MK-801 / Saline during ZT12–24). The groups were compared using Welch's *t*-test. It should be noted that the significance level of *P*-values is corrected using the Bonferroni correction. Error bar: SEM, $*P < 0.05$, $**P < 0.01$, $***P < 0.001$, n.s. = non significance.

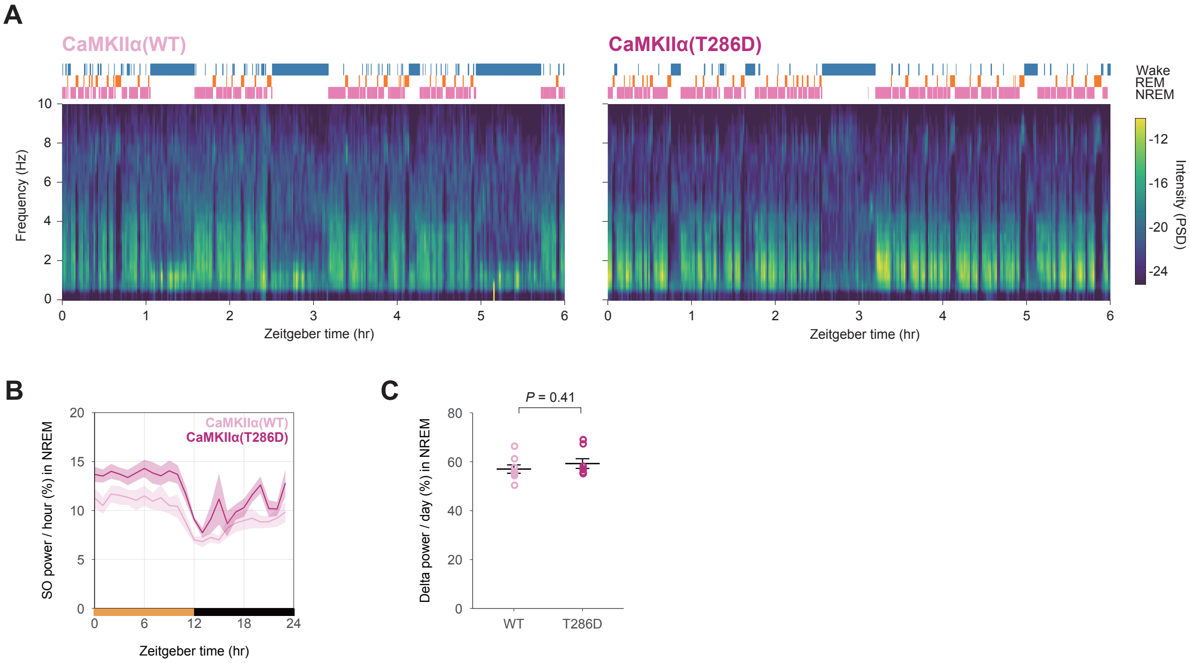


**fig. S10. Elevation of CaMKIIα kinase activity in E11 neurons facilitates NREM sleep quality.** (**A**) Representative sleep-wake features of E11-CaMKIIα (WT) (left) and E11-CaMKIIα (T286D) (right) mice are shown as hypnograms (top) and heat maps of EEG spectra (bottom). (**B**) Normalized delta (0.5–4 Hz) power during NREM sleep in E11-CaMKIIα (WT) and E11-CaMKIIα (T286D) mice ($n$ = 8 for each group). The groups were compared using Welch's $t$-test. (**C**) Hourly normalized slow-oscillation (SO) power (0.5–1Hz) during NREM sleep. Error bar: SEM.

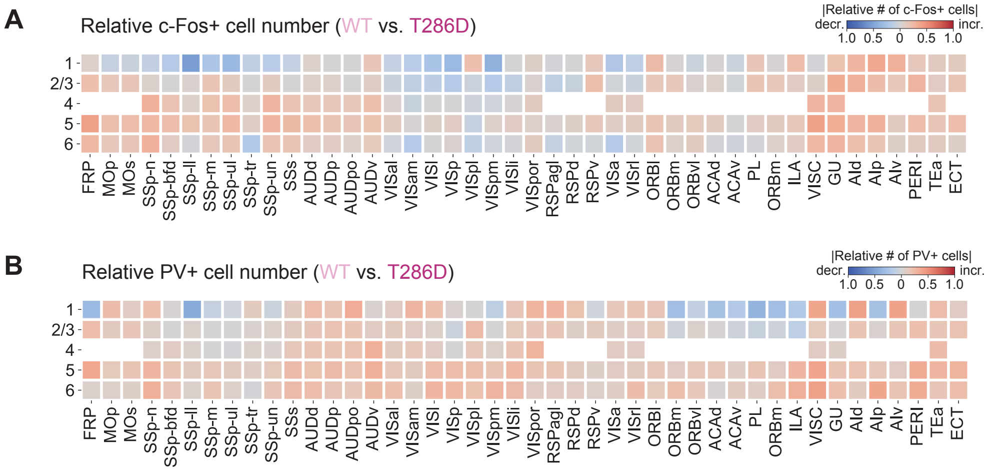


**fig. S11. Analysis of cortical c-Fos+ or PV+ cells in elevation of CaMKIIα kinase activity in E11 neurons.**

(**A** and **B**) The changes in c-Fos+ and PV+ cell number in the isocortex are represented as a heat map of relative cell number in each region. The E11-CaMKIIα (T286D) brains were compared to the E11-CaMKIIα (WT) brains, with red indicating an increase and blue indicating a decrease. The Allen Brain Atlas ontology is used to define brain region acronyms.

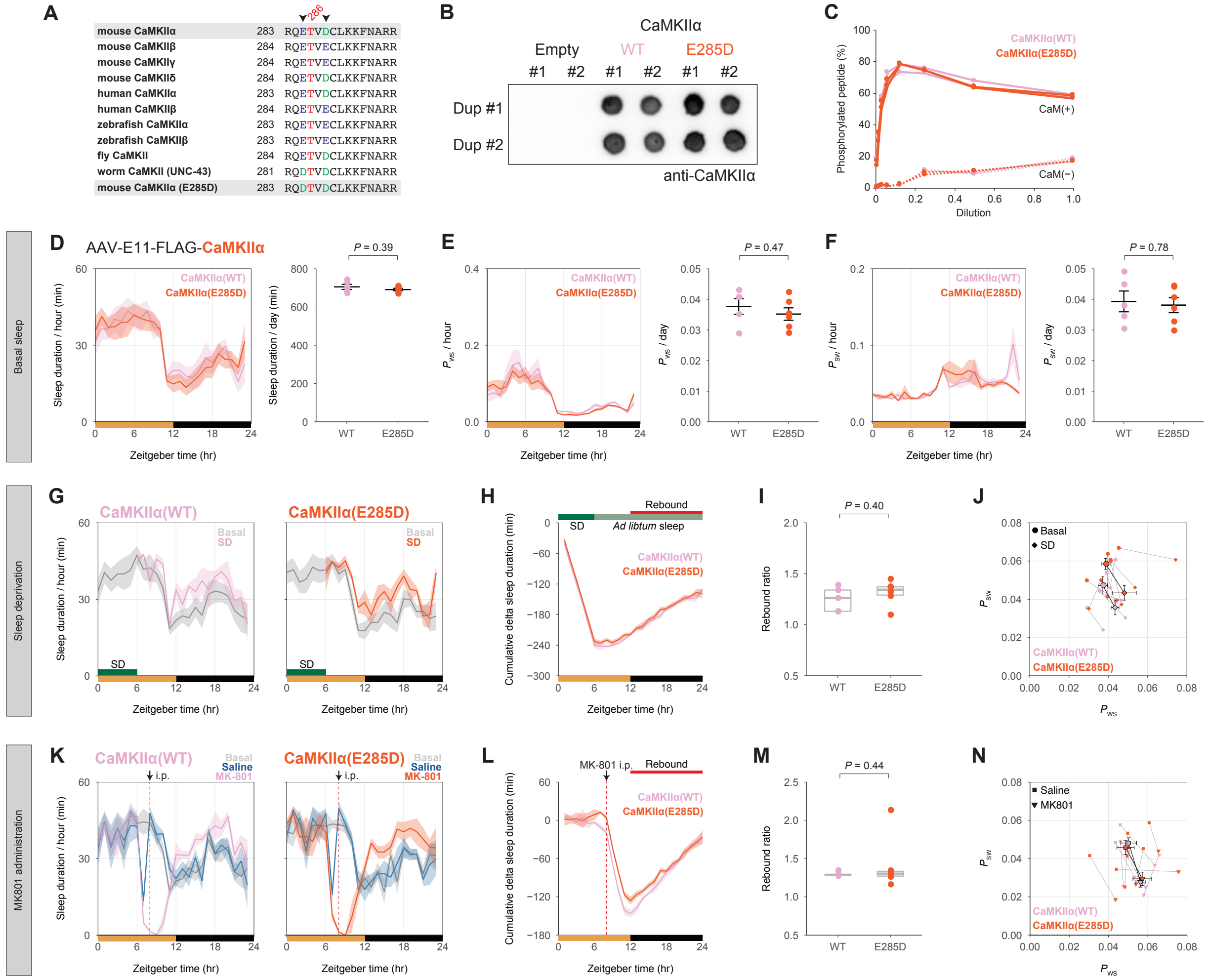


**fig. S12. Watermark mutation E285D in CaMKIIα does not affect the kinase activity, sleep architecture, and sleep homeostasis.**

(**A**) Sequence alignment of T286 (red) surrounding region of CaMKII family. The alignment was performed against the mouse (*M. musculus*) CaMKIIα/β/δ/γ, human (*H. sapiens*) CaMKIIα/β, zebrafish (*D. rerio*) CaMKIIα/β, fly (*D. melanogaster*) CaMKII, and worm (*C. elegans*) CaMKII analog (UNC-43). The residues of Glutamic acid (E) and aspartic acid (D) are colored in blue and green, respectively. The E285 and D288 positions of mouse CaMKIIα are indicated by black arrowheads. (**B**) Dot blots of HEK293T cell lysates transfected CaMKIIα expression vectors with CaMKIIα antibody ($n$ = 2 for each group). Vertical duplicates of each lysate were made. (**C**) CaMKIIα kinase activity assay with CaMKII peptide substrate. The phosphorylated peptide was calculated as a percentage of the total peptide ($n$ = 2 for each group). (**D**, **E**, and **F**) Daily sleep duration, $P_{\mathrm{WS}}$, and $P_{\mathrm{SW}}$ in E11-CaMKIIα (WT) ($n$ = 5, pink) and E11-CaMKIIα (E285D) ($n$ = 6, orange) mice. The groups were compared using Welch's $t$-test. (**G** and **H**) Hourly sleep duration and cumulative sleep loss (SD – Basal) in E11-CaMKIIα (WT) ($n$ = 5, pink) and E11-CaMKIIα (E285D) ($n$ = 6, orange) mice over 24 hours. Mice were allowed to behave freely for three days prior to SD, and the averaged value of sleep parameters is shown as "Basal." (**I**) Sleep rebound ratio (SD / Basal during ZT12–24). The groups were compared using Welch's $t$-test. (**J**) $P_{\mathrm{WS}}$ and

$P_{SW}$ scatter diagrams on Basal (circles) and SD (diamonds). Averaged values are represented by black-fringed dots, while individual mouse values are represented by other dots. Gray lines connect the dots of Basal and SD in each mouse. (**K** and **L**) Hourly sleep duration and cumulative sleep loss (MK-801 – Saline) in E11-CaMKIIα (WT) ($n$ = 5, pink) and E11-CaMKIIα (E285D) ($n$ = 6, orange) mice over 24 hours. The mice were allowed to behave freely for three days prior to the first injection, and the averaged value of sleep parameters is shown as "Basal." Each mouse received i.p. injection in turn: saline or 2 mg/kg MK-801 injection at ZT8, allowing at least 2 days between injections. (**M**) Sleep rebound ratio (MK-801 / Saline during ZT12–24). The groups were compared using Welch's *t*-test. (**N**) $P_{WS}$ and $P_{SW}$ scatter diagrams on Saline (squares) and MK-801 (triangles). Averaged values are represented by black-fringed dots, while individual mouse values are represented by other dots. Gray lines connect the dots of Saline and MK-801 in each mouse. Error bar: SEM, $^{*}P < 0.05$, $^{**}P < 0.01$, $^{***}P < 0.001$, n.s. = non significance.
